# Combining experiments and simulations to study the impact of multiple phosphorylations at the AT8 epitope of the Tau protein

**DOI:** 10.1101/2025.02.22.639620

**Authors:** Cynthia Lohberger, Jules Marien, Clarisse Bridot, Chantal Prévost, Diane Allegro, Mario Tatoni, Emmanuelle Boll, François-Xavier Cantrelle, Marlène Martinho, Valérie Belle, Caroline Smet-Nocca, Sophie Sacquin-Mora, Pascale Barbier

**Author notes:** Correspondence, INP, 27 bd Jean Moulin, 13005 MARSEILLE, IBPC, Université Paris Cité, CNRS, 75005, Paris, France. These authors contributed equally to this work (Cynthia Lohberger, Jules Marien).

## Abstract

Hyperphosphorylated Tau is a hallmark of Alzheimer’s disease leading to functional loss and fibrillar inclusions. We aim to understand how AT8 phosphorylation affects the Tau’s conformation and dynamics during early pathological transformation. We engineered an alanine Tau mutant to target specific phosphorylation sites restricted to the AT8 epitope, generating two distinct phosphorylation states. Using a combination of biophysical methods, and pCALVADOS forcefield, we found that AT8 phosphorylation do not alter the hydrodynamic radius or overall dynamics of Tau. Simulations revealed that local stiffening and extension at the AT8 epitope scale with phosphorylation extent. Interestingly, phosphorylation induces distant contact losses toward the N-terminus. This study advances our understanding of the structure-function-dynamics relationship of Tau in neurodegeneration.

## Introduction

Alzheimer’s disease (AD) is characterized by the presence of two key biomarkers: extracellular amyloid plaques formed by the accumulation of β-amyloid fibrils, and intracellular neurofibrillary tangles composed of aggregated hyperphosphorylated Tau protein, along with neurodegeneration [1]. Tau is an intrinsically disordered protein belonging to the microtubule-associated protein family [2]. The physiological role of Tau in stabilizing axonal microtubules, thereby ensuring proper neuronal activity, is mediated by its interaction with tubulin via the C-terminal MicroTubule-Binding Domain (MTBD) and a region within the Proline Rich Domain (PRD) [3]. While phosphorylation regulates this physiological function, it is also a critical factor in the pathological aggregation of Tau into neurofibrillary tangles.

In the central nervous system, Tau exists as six isoforms ranging from 352 to 441 amino acids, generated by differential splicing resulting in variations in the N-terminal domain (0N, 1N, 2N) combined with different numbers of MTBD repeats that are composed of three or four imperfectly repeated sequences of 18 residues (3R, 4R) [4]. Phosphorylation within the PRD has been shown to be involved in Alzheimer’s disease (AD) [5]. Building upon this clinical relevance, the present study focuses on pathological phosphorylations of Tau recognized by the AT8 antibody at Serine 202, Threonine 205, and Serine 208 [6]. We chose these specific phosphorylation sites as the AT8 antibody is commonly used in post- mortem diagnosis of Alzheimer’s disease and its staining on patients’ brains correlates with the different Braak stages defining the disease progression [7]. To isolate the effects of three and five specific phosphorylations including AT8 epitope, we used a Tau isoform in which 16 NMR-characterized phospho-sites were mutated to Alanine to abolish their phosphorylation by CDK2/cyclin A and GSK3β kinases. We define these phosphorylation states as p-state 1 (Serine 202, Threonine 205, and 212 are phosphorylated: pS202 + pT205 + pT212) and p-state 2 (p-state 1 with additional phosphorylation on Serine 198 and 208; pS198 + pS202 + pT205 + pS208 + pT212) (Fig 1 and S1) and compare them to the unphosphorylated AT8 mutant. Previous NMR structural studies have revealed that pS202/pT205/pT212 in p-state 1 induces a local conformational change characterized by a kink at residue G207. Interestingly, this kink is disrupted by the additional phosphorylation of Serine 208 [8,9]. Furthermore, while pS202/pT205 is unable to form oligomers, the triple phosphorylation pS202/pT205/pS208 exhibits a propensity to form pathological oligomers [8,9]. These findings suggest that the phosphorylation of Serine 208 plays a critical role in promoting detachment from microtubules and Tau aggregation, thereby contributing to the pathogenesis of neurodegenerative diseases by local conformational modification [10]. To assess the impact of this local conformational change on the overall conformation and dynamics of Tau, we combined experimental and simulation studies to determine the effect of multiple and specific phosphorylation on the local and global dynamics of Tau. After the control of the specific phosphorylation sites by NMR studies, hydrodynamic radii were measured by Dynamic Light Scattering (DLS) and Analytical UltraCentrifugation (AUC). Additionally, Site-Directed Spin Labeling combined with EPR spectroscopy was used to probe the local dynamics around the AT8 epitope comparing unphosphorylated AT8 mutant to p-state 1 and p-state 2. Furthermore, Molecular Dynamics (MD) simulations have been proved to be a method of choice to study the impact of mutations or post-translational modifications on the conformational dynamics of proteins. However, simulating long IDPs via MD has proven to be a significant challenge due to their rugged conformational space and their large fluctuations in size. Coarse-grained models are promising attempts to generate Conformational Ensembles (CoE) at low computational cost compared to all-atom simulations. A popular type of force fields are ones based on the HydroPhobicity Scale (HPS) model by Kapcha and Rossky [11], among which the machine-learned CALVADOS FF is becoming a staple in the field [12]. However, the phosphoparameters by Rauh et al. [13] did not yet exist when performing the simulations of the present article, we thus assembled a new forcefield named pCALVADOS from Perdikari et al. phosphoparameters [14] and performed simulations matching the experimental sequences.

**Figure 1:**
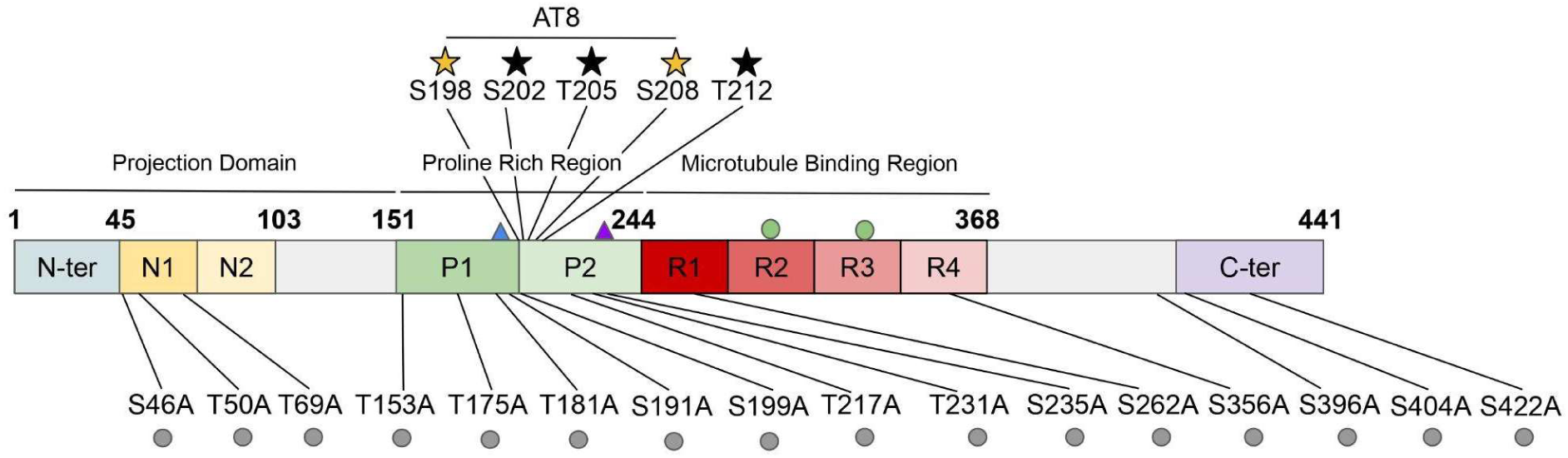
Domain organization and phosphorylation sites of wild-type longest human Tau isoform and its mutants designed to investigate the role of specific phosphorylation events in Alzheimer’s disease. The gray circles represent the 16 Serines/Threonines mutated into Alanine. p-state 1: Serine 202 and Threonines 205 and 212 are phosphorylated indicated by black stars. p-state 2: Serines 198 and 208 are additionally phosphorylated compared to p-state 1 indicated by yellow stars. Additional mutations for Site-Directed Spin Labeling EPR experiments are indicated: green dots for the two natural cysteines (C291 and C322) mutated in Alanine and triangles the introduced cysteines: (S185C, blue) and (A227C, purple).

The computational assessment of the local dynamics of an IDP is a complex task as standard alignment-based methods such as RMSD and RMSF lose all meaning in disordered systems. We thus employed new metrics developed by Marien et al. [15] and specifically tailored for IDPs and IDRs (Proteic Menger Curvature, Local Curvature and Local Flexibility) [16].

Altogether, all data corroborated the absence of impact of the different phosphorylation states on the global shape and dynamics of Tau. Nevertheless, simulations allowed us to identify local stiffening and extension increasing with the phosphorylation states. Interestingly, phosphorylation goes beyond the immediate AT8 region, revealing distant contact losses, directed towards the N-terminus, potentially influencing Tau’s function. These findings suggest a potential mechanism in which specific phosphorylation could modulate Tau’s interactions and function, contributing to neurodegeneration. Combining experimental studies with the pCALVADOS model, we provided a robust approach for future studies on Tau’s structure-function-dynamics and more generally for studying intrinsically disordered proteins and the impact of post-translational modifications on their conformational ensembles.

## Results

### Characterization of phosphorylated and unphosphorylated AT8 mutants

To study the specific effects of Tau phosphorylation at the AT8 epitope, a Tau mutant was generated in which 16 Ser/Thr residues targeted by CDK2/cyclin A and GSK3*β* kinases were replaced by alanine, while preserving the phosphorylation sites recognized by the AT8 antibody (Figs 1 and S1). The mutant is then converted into two distinct phosphorylation states using sequential kinase treatment. P-state 1 is obtained by incubation with CDK2/cyclin A kinase, whereas p-state 2 was generated by subsequent incubation of p-state 1 with GSK3*β*.

The overall phosphorylation levels are first assessed by SDS-PAGE and MALDI-TOF mass spectrometry (Fig S2, Table S1). MALDI-TOF measurements confirmed the expected mass decrease of the AT8 mutant relative to WT Tau and revealed progressive mass increases upon phosphorylation. However, because intact protein MALDI-TOF spectra of Tau (∼46 kDa) exhibit a relatively low resolution (FWHM ≈ 300- 500 Da), individual phospho-species differing by a single phosphorylation event (∼80 Da) cannot be resolved. Consequently, the observed masses correspond to the average mass of heterogeneous phospho-proteoform populations (in addition to Na+/K+ adducts and isotopic heterogeneity) rather than to single phosphorylation states. Site-specific phosphorylation patterns and phosphorylation levels were therefore determined primarily by high-resolution NMR spectroscopy.

Given the high sensitivity of the N-H amide resonances to modifications of their chemical environment such as phosphorylation events, chemical shift perturbations are detected in the 2D ^1^H-^15^N HSQC spectra for Ser/Thr phosphorylation sites and neighboring residues (Fig 2). In IDPs, Ser/Thr phosphorylation typically gives rise to resonances in otherwise sparsely populated regions of the spectrum, facilitating their identification. In addition, integration of the respective amide resonances of the phosphorylated and non-phosphorylated forms of a given Ser/Thr residue allows site- specific quantification of phosphorylation levels.

**Figure 2:**
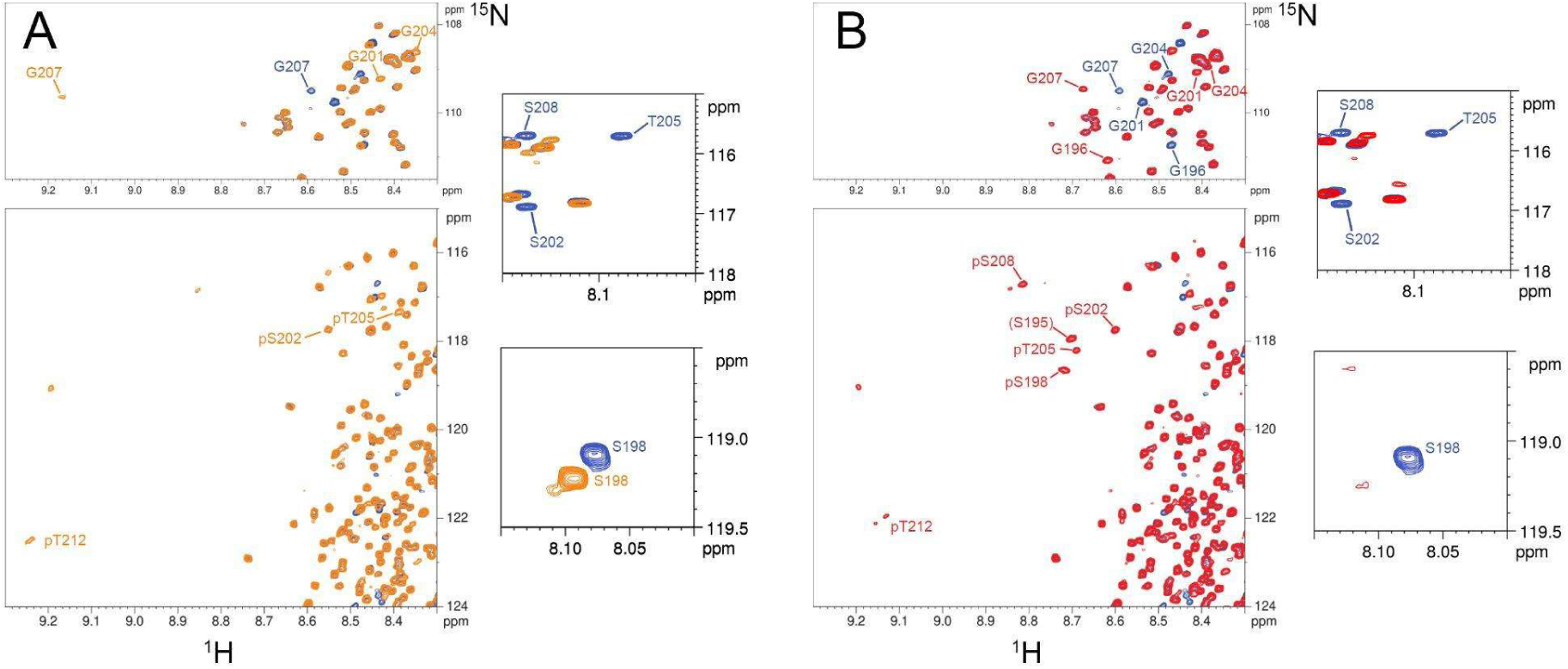
Expanded views of NMR ^1^H-^15^N HSQC spectra of the unphosphorylated AT8 mutant (blue), the mutant in p-state 1 (orange) and in p-state 2 (red), highlighting the site-specific phosphorylation sites in each phosphorylation state. (A) Comparison of the spectra of the mutant in p-state 1 and the unphosphorylated state. (B) Comparison of the spectra of the mutant in p-state 2 and the unphosphorylated state. The regions of phosphorylated Ser/Thr and Gly (left panels) are depicted to show the chemical shift perturbations of resonances upon phosphorylation. S202/T205 double phosphorylation in p-state 1 and S202/T205/S208 triple phosphorylation in p-state 2 have a differential impact on G207 resonance which is found in the center of the AT8 motif. The unphosphorylated S202, T205, S208 and S198 resonances are depicted (right panels) to show their disappearance or shift upon phosphorylation.

The ^1^H-^15^N HSQC spectrum of the Tau AT8 mutant in its non-phosphorylated state has been assigned using a set of three-dimensional backbone assignment experiments. Additional 3D NMR experiments on both phosphorylated Tau AT8 mutants enabled the assignment of resonances for p-state 1 and p-state 2 that have been shifted upon phosphorylation. By interpreting the NMR spectra of the Tau mutant in its non-phosphorylated or phosphorylated states, in combination with mass spectrometry analyses (Figs S2 and S3), we concluded that p-state 1 results in quantitative phosphorylation of S202, T205 and T212. This conclusion is supported by the appearance of new resonances assigned to the phosphorylated forms of these residues together with the disappearance of the corresponding non-phosphorylated resonances. Phosphorylation also induces significant perturbations in neighboring residues, including S198 and S208. In addition, a pronounced chemical shift change was observed for G207, consistent with the formation of a local β-turn induced by double phosphorylation at S202 and T205, as previously reported [8,9]. The NMR spectra of p-state 1 also revealed two additional phospho-resonances that could not be unambiguously assigned (Figs S2 and S3). These signals indicate the presence of additional CDK2-dependent phosphorylation events outside the AT8 epitope, although their precise locations and occupancies could not be determined. The presence of such minor phospho-species is consistent with the larger-than-expected mass increment observed by MALDI-TOF analysis.

In p-state 2, two additional phosphorylation events were detected on S198 and S208 both reaching phosphorylation levels close to 100% (Figs 2 and S3). In contrast to the double phosphorylation of S202 and T205 in p-state 1, triple phosphorylation pattern S202/T205/S208 in p-state 2 induces a much smaller perturbation of the G207 resonance relative to the non-phosphorylated state. This observation is consistent with previous reports suggesting that phosphorylation of S208 disrupts the *β-*turn conformation stabilized by phosphorylation of S202/T205, thereby favoring a more extended local conformation [9].

Thus, NMR analysis demonstrates that the two engineered phospho-states are dominated by the intended phosphorylation patterns. These two well-defined phosphorylation states therefore provide a suitable framework to investigate the effects of distinct phosphorylation patterns within the AT8 epitope while minimizing interference from phosphorylation events elsewhere in the protein.

### Determination of the hydrodynamic radius by different methods

To investigate the hydrodynamic properties of Tau and of the unphosphorylated and phosphorylated AT8 mutants, we performed Dynamic Light Scattering (DLS) and Analytical UltraCentrifugation (AUC) measurements. In DLS experiments, correlograms show a typical decay pattern expected for a diffusing species (Fig S4). The correlation coefficient starts high (0.9), indicating strong initial correlation, and decays smoothly towards zero as time increases. This decay is consistent with Brownian motion of particles in solution and high quality of the provided data. Although DLS is primarily used for globular proteins, it can still provide information about conformational ensembles in IDPs and especially aggregation states. The volume- weighted distribution showed one peak representing more than 80% of species present in solution. Because DLS measures an apparent hydrodynamic radius that is generally concentration-dependent due to interparticle interaction in non-ideal solution, we then plotted the apparent hydrodynamic radius (Rh) as a function of protein concentrations and extrapolated with a linear regression to infinite dilution (Fig 3, Table S2) to best estimate the true Rh (Rh_0_). The confidence interval of the intercepts corresponding to the Rh of each Tau sample was calculated as described in materials and methods. WT Tau exhibited a Rh_0_ of 5.0 nm with a confidence interval of 1.3 nm in agreement with this obtained previously [17,18]. The unphosphorylated AT8 mutant displayed the same value of Rh_0_ of 5.0 ± 1.2 nm. The observation that the mutant exhibits a hydrodynamic radius comparable to that of wild-type Tau suggests that this mutant provides a suitable model for investigating AT8-specific phosphorylation. The p-state 1 has a Rh_0_ value of 5.8 ± 1.2 nm and the p-state 2 displays a Rh_0_ of 5.0 ± 1.8 nm. The confidence intervals for the four different samples overlap, indicating no significant difference in the hydrodynamic radii. By consequence, the phosphorylation state of AT8 Tau mutant has no effect on the global shape of this protein. On the other hand, the positive slope of the linear regression indicates an increase of Rh as a function of the concentration. This tendency is strong for both the unphosphorylated AT8 mutant (Fig 3B) and the p-state 2 (Fig 3D). In contrast, WT Tau (Fig 3A) and the p-state 1 phosphorylated mutant (Fig 3C) exhibit a much lower concentration dependence due to non-ideality of the solution.

**Figure 3:**
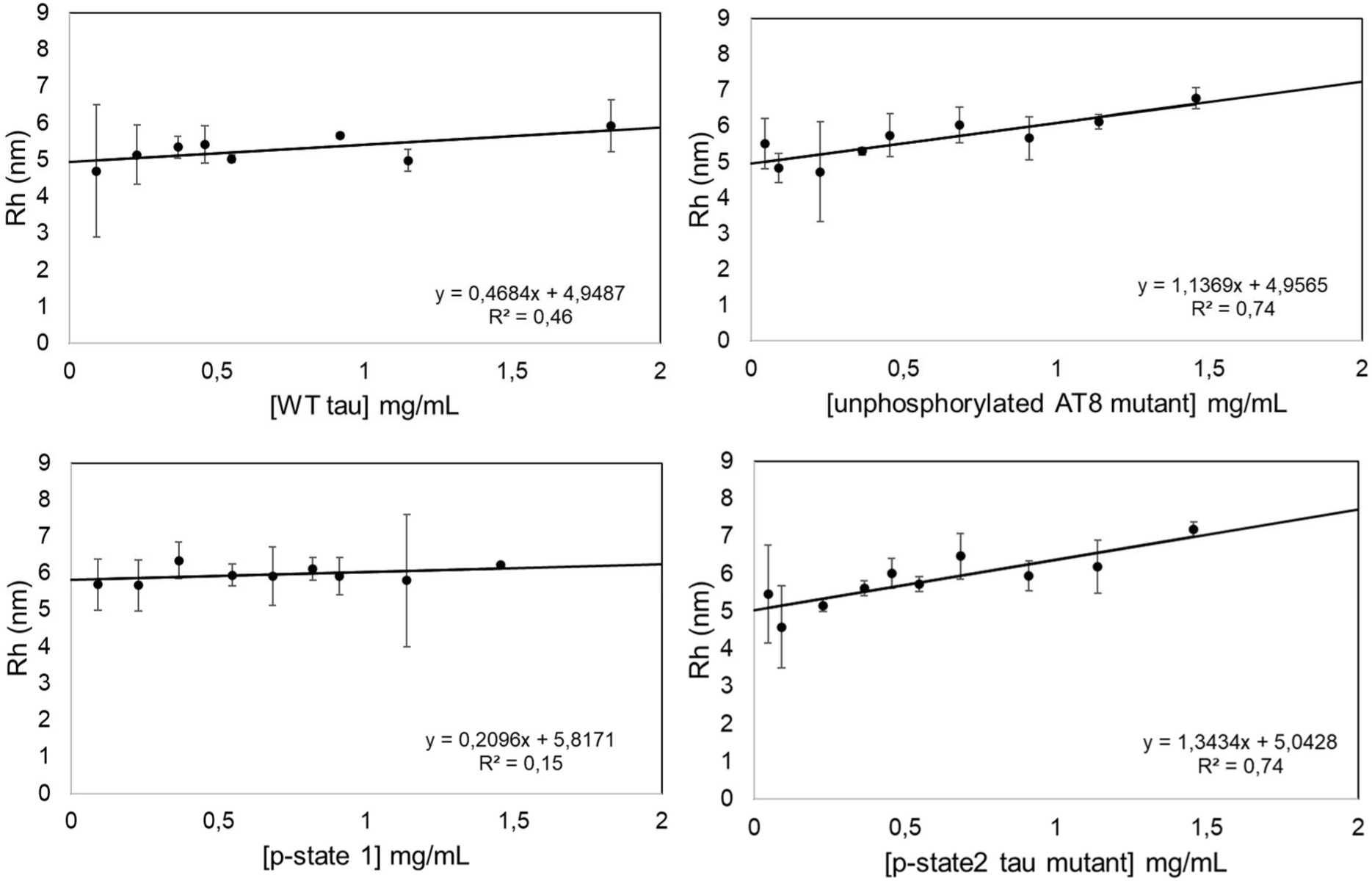
Hydrodynamic radius of WT Tau, unphosphorylated AT8 mutant, p-state 1 mutant and p-state 2 mutant as a function of protein concentration. Apparent Rh measured from DLS experiments and the linear regression in black. (A) WT Tau (concentration range 0.09-1.83 mg/mL). (B) unphosphorylated AT8 mutant (concentration range 0.045-1.45 mg/mL). (C) Tau mutant in p-state 1 (pS202 + pT205 + pT212) (concentration range 0.09-1.45 mg/mL). (D) Tau mutant in p-state 2 (pS198 + pS202 + pT205 + pS208 + pT212) (concentration range 0.045-1.45 mg/mL). The error bars indicate the standard deviation of the average Rh value obtained from a triplicate. The Rh_0_ was obtained by extrapolating the linear regression to infinite dilution and its confidence interval was calculated as described in the statistical section of materials and methods.

A second experimental approach was employed to confirm the absence of impact of phosphorylation on the global conformation of Tau. The sedimentation velocity experiment in Analytical UltraCentrifuge is a highly reproducible and quantitative method that allows the determination of the sedimentation coefficient (S). This parameter reflects both the molecular mass and the hydrodynamic shape of the protein. The displacement of the protein by the centrifuge force, measured by changes in absorbance as a function of the centrifugation radius over time, is analyzed as a distribution in which each peak corresponds to a different molecular species with a distinct sedimentation coefficient.

For WT Tau, the C(S) distribution shows a major peak (90%) at a S_app_ equal to 2.5 ± 0.6 Svedberg (S), corresponding to the monomeric form of Tau (Fig 4A). A shoulder observed around 4 S suggests the presence of a small amount (10%) of a different Tau species. In the case of the unphosphorylated AT8 mutant (Fig 4B), the main peak (82%) corresponding to monomeric Tau is centered at S_app_ equal to 2.4 ± 0.1 S. This peak appears sharper and narrower compared to that of WT Tau, suggesting a reduced conformational variability within the sample. As for WT Tau, we observed 3% of a second species at 4S. Additionally, a smaller peak (15%) appears at 1.5 ± 0.1 S. For the AT8 mutant in p-state 1 (Fig 4C), the C(S) distribution also displays three peaks as for the unphosphorylated mutant: a major one (89%) corresponding to the monomeric Tau at 2.5 ± 0.3 S, another peak at 4S (2%) and a smaller one at 1.5 S (9%). Finally, the AT8 mutant in p-state 2 (Fig 4D) showed the same sedimentation profile with one peak at 2.4 ± 0.3 S corresponding to the monomer (85%), a second at 4S (5.5%) and a third one at approximately 1.5 S (9.5%). Due to hydrodynamic non-ideality, which increases intermolecular friction through crowding effects, the apparent sedimentation coefficients S_app_ obtained for the monomeric form of Tau of each sample were measured, corrected to standard conditions (20°C in water; S_20,W_) plotted for various Tau concentrations, and analyzed by linear regression. This analysis yielded the standard sedimentation coefficient at infinite dilution (S^0^_W,20_) representing the intrinsic sedimentation behavior of the molecule in the absence of interparticle interactions (Fig 5). We obtained a sedimentation coefficient S^0^_W,20_ of 2.4 ± 0.3 S, 2.4 ± 0.3 S, 2.2 ± 0.5 S and 2.4 ± 0.3 S for WT Tau, unphosphorylated AT8 mutant, mutant in p-state 1 and p-state 2 respectively (intercept ± confidence interval). The confidence intervals calculated as described in Materials and Methods section overlapped for the four different Tau monomers indicating no significant difference in the sedimentation coefficients. This points towards a lack of effect of the mutations and of the phosphorylation state of Tau in monomeric state.

**Figure 4:**
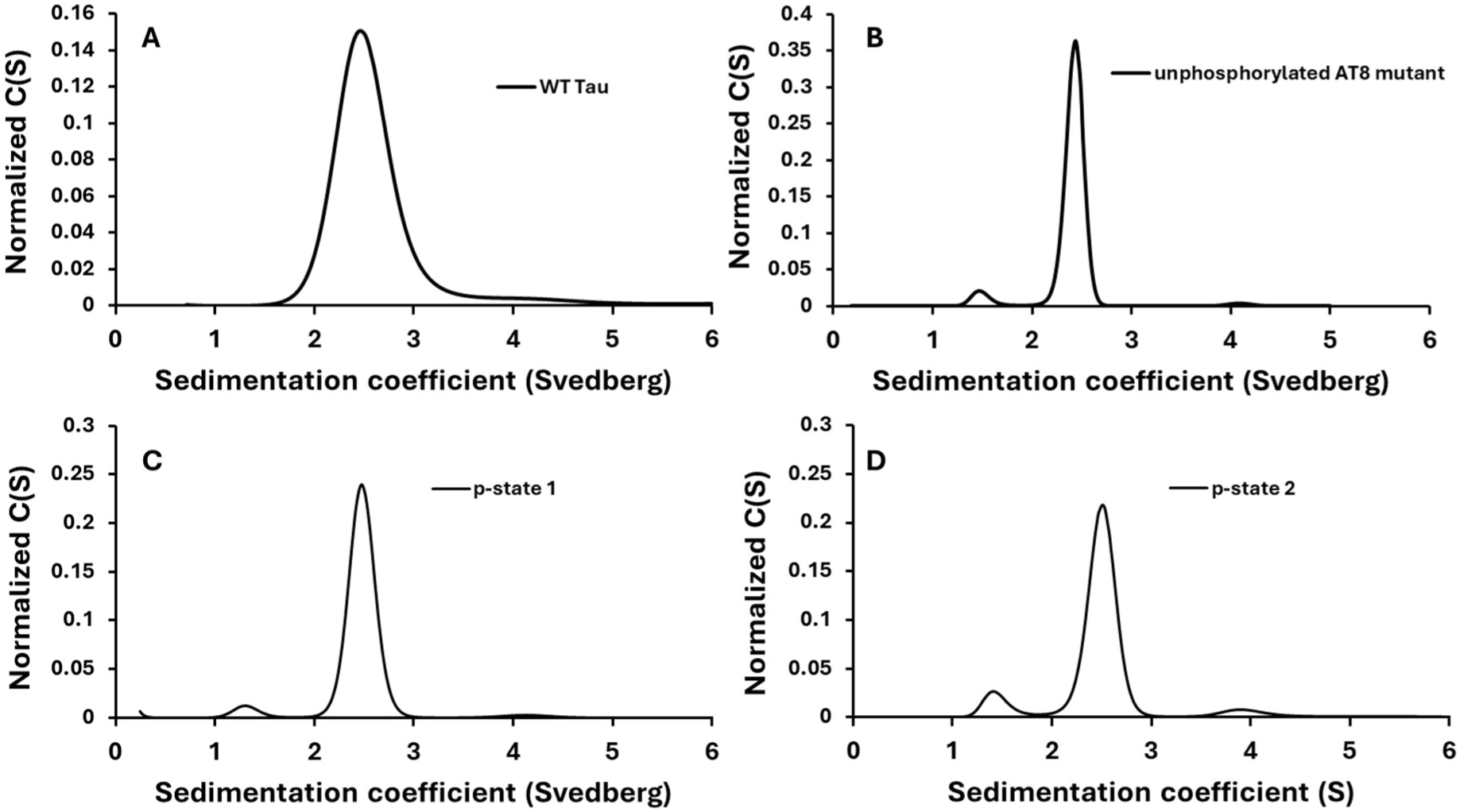
Analytical sedimentation velocity experiment analysis of WT Tau and its AT8 mutant unphosphorylated and phosphorylated in p-state 1 and 2. Normalized Sedimentation coefficient distribution determined by Sedfit analysis of WT Tau **(A)**, unphosphorylated AT8 mutant **(B)**, AT8 mutant in p-state 1 **(C)** and AT8 mutant in p-state 2 (D).

**Figure 5:**
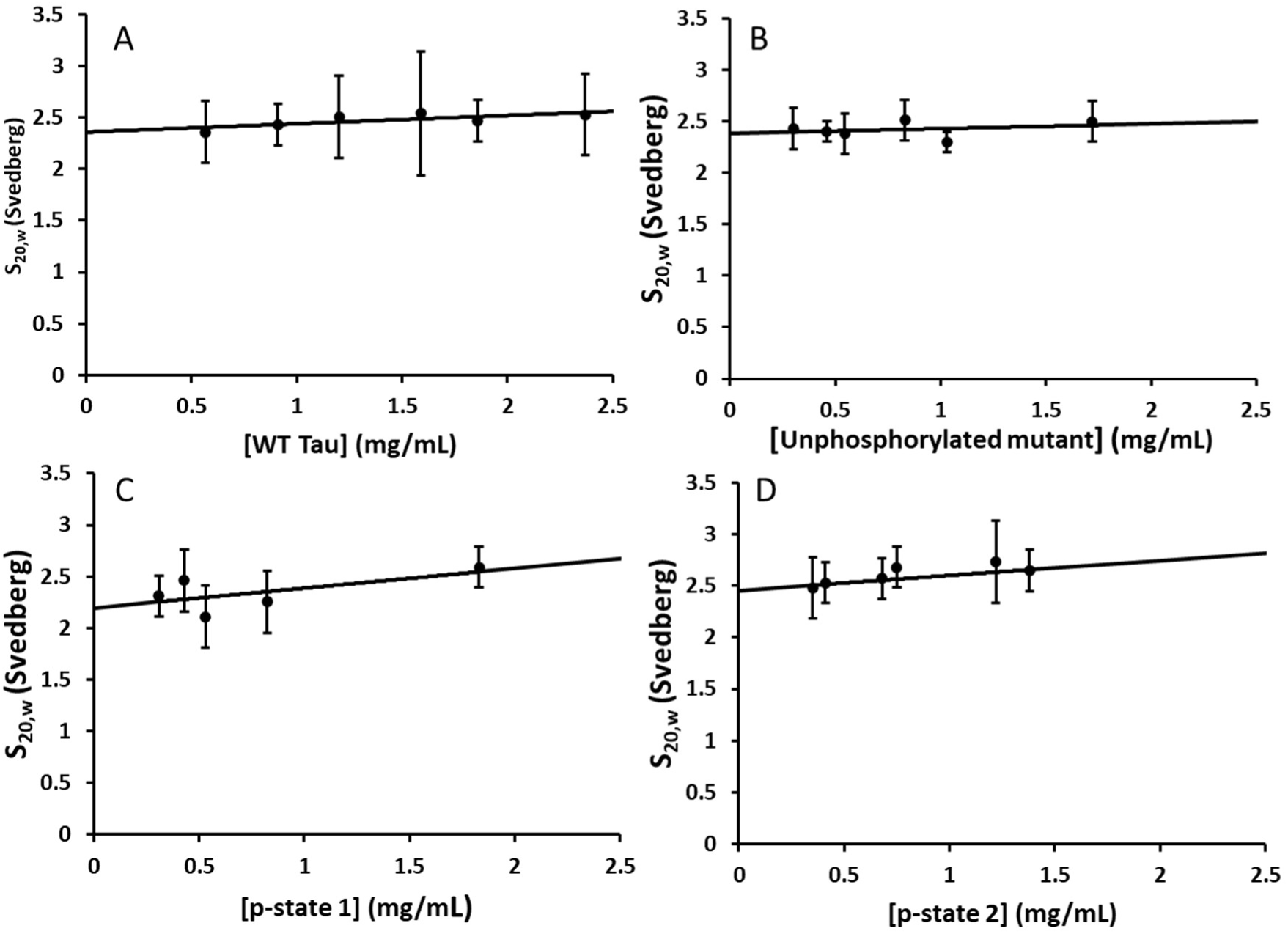
Determination of standard sedimentation coefficient S^0^_W,20_ by sedimentation velocity experiments. The apparent sedimentation coefficients (S_app_) of WT tau **(A)**, unphosphorylated AT8 tau mutant **(B),** and its p-state 1 **(C)** and p-state 2 **(D)** phosphorylated forms determined from figure 4 at several concentrations were corrected to standard conditions (20°C, water as solvent) to yield S_20,W_ values. These values were plotted against protein concentration, and a linear fit was applied to account for concentration-dependent effects due to hydrodynamic non-ideality. The extrapolation of this fit to infinite dilution yields the standard sedimentation coefficient, S^0^_20,W_, representing the intrinsic sedimentation behavior of the molecule in the absence of interparticle interactions. Its confidence interval was calculated as described in the statistical section of Materials and Methods.

Values of hydrodynamic radii were estimated from the sedimentation coefficient measured in sedimentation velocity experiments with the Svedberg equation (see Materials and Methods). The obtained values are in accordance with those measured directly by DLS and confirm the absence of effect of the mutations and of phosphorylations on the global conformation of Tau (Table1). Values for the hydrodynamic radius (Rh_0_) were computed using the Nygaard method specifically developed with IDPs in mind. According to Pesce et al, this method is highly suitable for determining the Rh_0_ of Tau [17]. Consistent with this, we obtained a computed Rh_0_ value of 5.2 +/- 0.7 nm. The hydrodynamic radii obtained using both experimental and computed methods are consistent within this study and others [17], suggesting a good reliability of the results (table 1).

**Table 1:** Hydrodynamic radius of wild-type and phosphorylated Tau variants as determined by, DLS, and AUC experiments and pCALVADOS simulations.

| Protein | DLS experimental<br>$Rh_0$ (nm) | AUC experimental<br>$Rh_0$ (nm) | Nygaard method<br>$Rh_0$ (nm) |
| --- | --- | --- | --- |
| WT Tau | $5.0 \pm 1.3$ nm | $4.5 \pm 0.6$ nm | $5.2 \pm 0.7$ nm |
| unphosphorylated AT8 mutant | $5.0 \pm 1.2$ nm | $4.5 \pm 0.6$ nm | $5.2 \pm 0.7$ nm |
| AT8 Tau mutant p-state 1 (pS202, pT205, pT212) | $5.8 \pm 1.2$ nm | $4.6 \pm 1.1$ nm | $5.1 \pm 0.7$ nm |
| AT8 Tau mutant p-state 2 (pS198, pS202, pT205, pS208, pT212) | $5.0 \pm 1.8$ nm | $4.5 \pm 0.6$ nm | $5.1 \pm 0.7$ nm |

### Local dynamics

DLS and AUC experiments revealed no significant differences between p-states in the global hydrodynamics of Tau. We therefore asked whether a change could be observed in the local dynamics, at the residue scale. The transient nature of intramolecular interactions can be investigated via the simulated trajectories using contact maps derived from the simulated CoEs (Figs S5, S6 and S7 and Materials and Methods). All contacts made between two residues in all systems are present less than 10% of the simulated time, indicating a very dynamic behavior of the peptide chain (Fig S6). To assess the local impact of the mutations and of the phosphorylations, we decided to use the WT Tau simulation as a reference. We subtracted the contact map of the wild-type to the mutant and its different p-states to highlight which contacts are impacted. Fig 6A displays the difference between the unphosphorylated AT8 mutant and WT Tau. Long-range interactions are barely modified by the alanine mutations with differences compared to reference of less than 1%, and local interactions (within five residues) mostly show negligible losses of contacts inferior to 5%. This similarity of contact profiles between the unphosphorylated mutant and the wild type confirms that the former can be considered a reliable approximation of the latter, and that both can be equally considered for the comparison with the phosphorylated states.

**Figure 6:**
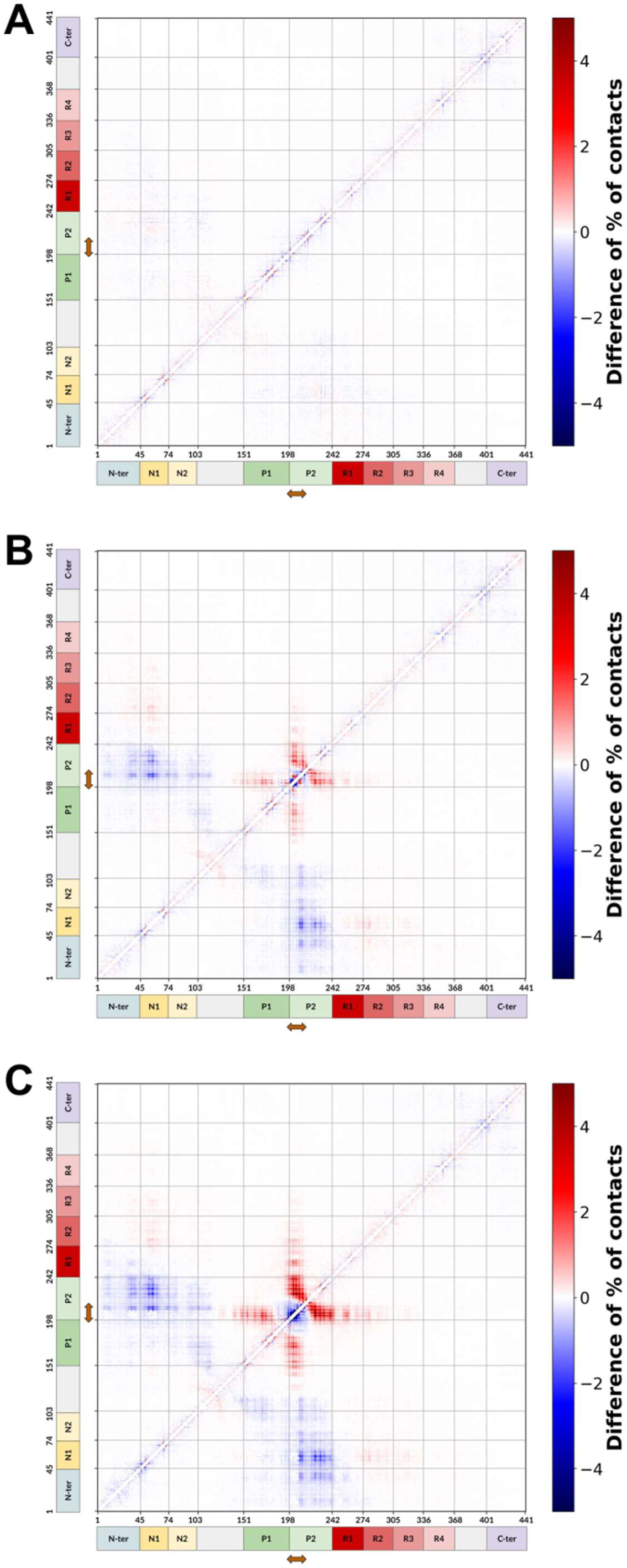
Difference of percentages of contacts per residue pairs between the unphosphorylated Tau wild type simulation and the AT8 mutant (A), the mutant in p-state 1 (B), the mutant in p-state 2 (C). Contact gains are in red, contact losses are in blue. The AT8 epitope is indicated by a brown arrow.

This comparison with the phosphorylated sequences reveals interesting local and long-range effects. Contacts are strengthened within 50 residues of the epitope, notably with R170, K174, K190 and R194 on the N-terminal side and with R221, K224, K225, R230, K234, K240 and R242 on the C-terminal side (Figs 6B and 6C). On the other hand, and quite surprisingly, contacts are weakened at very long residue-range (up to 200 amino-acids away) when the epitope is phosphorylated, with a stronger effect on residues E53, D54, E55, E57, E58 and a weaker one for D34, E35, E38, E73, E104 and E105. The effect is also only present for the N-terminal part of Tau, which indicates that the unphosphorylated epitope has a preferential affinity with it. These contact differences also increased as the number of phospho-residues increased, suggesting that they are of electrostatic nature.

### Local curvatures and flexibilities

In order to decipher changes in backbone dynamics, we employed computational metrics specifically designed to make sense of the backbone CoE of IDPs without the need for alignments or reference structures. The Local Curvatures (LCs) and Local Flexibilities (LFs), derived from the Proteic Menger Curvatures (PMCs) distributions of the residues, can assess the average curvature of the backbone and its fluctuations [16]. We previously introduced these metrics to characterize bending of the protein backbone induced by counterion bridges in multi-phosphorylated peptides [15]. Hence, we once again used the WT Tau simulation as a reference to assess the effects of the mutations and phosphorylations on the local curvature and flexibility of the backbone. The lack of significant differences in LCs and LFs at the substitution sites (Fig S5 and S7) confirms that the Tau mutant serves as an accurate structural surrogate. This validates our model, ensuring that observed changes are attributable to phosphorylation rather than the mutations themselves. Notably, phosphorylation triggers a pronounced effect on backbone dynamics spanning residues 195 to 215 (Fig 7). Local stiffening and elongation of the backbone, probably due to electrostatic repulsion from the addition of multiple phosphorylations, is observed, as both the curvature and the flexibility of the epitope residues are reduced. This effect depends on the number of phosphorylation as LCs are only decreased by -0.025 Å⁻¹ for P203 and G204 for p-state 1, whereas most residues from A199 to P206 see their LC drop by more than -0.025 Å⁻¹ for p-state 2 (Fig 7A). The difference between p-states is even more noticeable with LFs as differences with the wild-type for P200 and S210 are three times higher for p-state 2 than for p-state 1 (Fig 7B).

**Figure 7:**
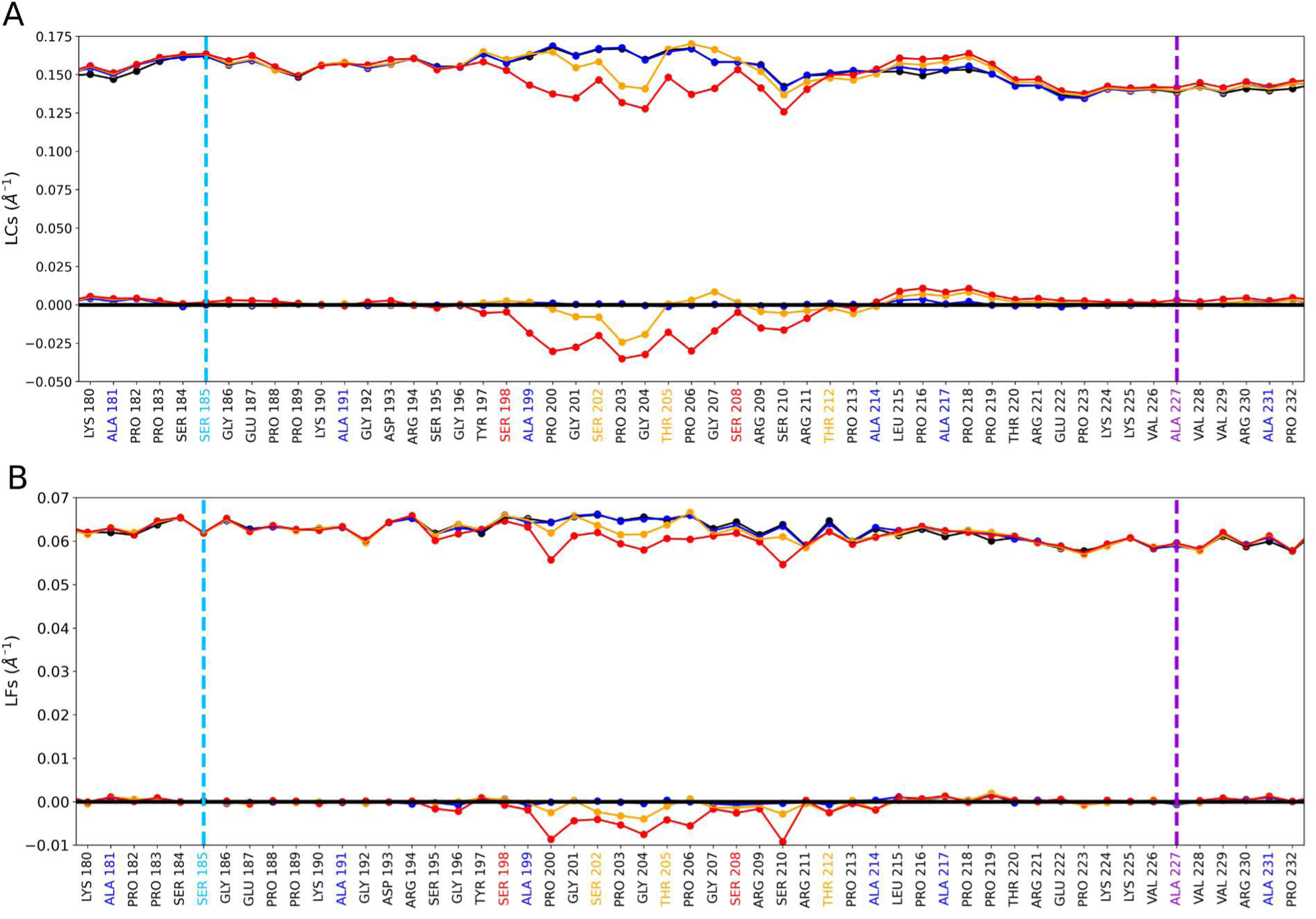
Local Curvatures (A) and Local Flexibilities (B) of the AT8 epitope. Upper curves on each graph represent Local Curvatures (A) and Local Flexibilities (B) for WT Tau (black), the unphosphorylated AT8 mutant (blue), the mutant in p-state 1 (orange) and the mutant in p-state 2 (red). The lower curves on each graph represent the difference regarding the wild type. Positive values represent a gain in curvature/flexibility respectively and negative values a loss. Residues in blue have been mutated to alanine compared to Tau wild type. Residues in orange are phosphorylated in p-state 1, residues in red in p-states 1 and 2. The mutated sites used in the EPR experiments for S185C and A227C are indicated in blue and purple dashed lines respectively.

Spin-labeling combined with EPR spectroscopy was used to experimentally assess the local flexibility at two specific positions (185 and 227) where single cysteine residues were introduced to graft nitroxide spin labels. These cysteine variants were available from a previous study focused on PRE and PRI NMR experiments [19]. This technique is very powerful to detect small changes in local flexibility of the protein backbone and particularly sensitive to report local structural changes in IDPs [20,21]. For each position, the EPR spectra showed very narrow lines indicative of high flexibility as expected for an intrinsically disordered region (Fig S8). No significant difference was observed after phosphorylation at p-state 1 and p-state 2. This experimental observation reinforces the findings brought by the calculations of LCs and LFs that showed modifications restricted to the close AT8 region without any fluctuation of these parameters neither downstream nor upstream of this region.

To summarize, pCALVADOS simulations of the backbone dynamics and derived hydrodynamic radii agree with the experimental data. In both cases we demonstrated that there is no effect on the global conformation of Tau. More interestingly, we show a reduction of the curvature and flexibility of the backbone around the AT8 epitope that could hint towards a role of phosphorylation for Tau dynamics.

## Discussion

Protein phosphorylation is a crucial posttranslational modification catalyzed by kinases which transfer phosphate groups from ATP to specific amino acid residues. This process regulates a wide range of cellular functions, including cell signaling, proliferation, apoptosis, and maintenance of neuronal function [22]. Among these enzymes, the Serine/Threonine kinases constitute an important part of the human kinome, targeting the hydroxyl groups of Serine or Threonine residues. These kinases also phosphorylate the Tau protein, thereby modulating its physiological and pathological roles. In neurodegenerative diseases such as Alzheimer’s, the equilibrium between kinases and phosphatases (enzymes that remove phosphorylation) is disrupted, resulting in Tau hyperphosphorylation. Several phosphorylation sites of Tau have been identified as pathological. Among them, phosphorylation at Serine 202, Threonine 205, and Serine 208 defines the AT8 epitope, recognized by a specific antibody routinely employed in the post-mortem diagnosis of Alzheimer’s disease [6] and which the intensity staining in brain tissue correlates with the Braak stages that characterize the progression of the disease [7]. The longest isoform of human Tau has 85 potential phosphorylation sites along its 441 amino acid sequence, 40 of which have been characterized experimentally [23,24]. Because understanding their role presents an analytical challenge, previous studies predominantly used phosphorylated Tau peptides, which failed to recapitulate the conformational and functional properties of the complete protein. By consequence in this study, we aimed to restrain the phosphorylation sites within the full-length Tau protein. To achieve this, specific residues previously identified by NMR as phosphorylation sites of CDK2/cyclin A and GSK3β kinases *in vitro* [8,9] were substituted with Alanine to prevent phosphorylation [22,23], except for the pathological epitope recognized by the AT8 antibody. Moreover, IDPs, such as Tau, lack a stable 3D structure, making them challenging to study their conformation in solution and giving them significant aggregative properties. We used DLS and AUC to determine the global shape of WT Tau, the unphosphorylated mutant, p-state 1 and p-state 2 through the sedimentation coefficient and the hydrodynamic radius parameters determination. The hydrodynamic radius of WT Tau obtained by DLS analysis in our study is consistent with values previously reported using the same methodology [25,26]. Unexpectedly, the addition of phosphorylation on the AT8 epitope has no effect on the Tau hydrodynamic radius. Because in DLS, the presence of a small number of larger particles contributes significantly to light scattering, we completed our study using Analytical UltraCentrifugation (AUC) sedimentation velocity (SV) experiments. Indeed, AUC is a very powerful technique to characterize macromolecules or mixtures of macromolecules in solution by separating them using centrifuge force, allowing the measurement of their hydrodynamic properties: their sedimentation coefficient S [27] and their hydrodynamic radius (see Materials and Methods section). The primary advantage of this method lies in its versatility, as it is applicable to spherical proteins, elongated structures, and IDPs, enabling the identification and quantification of distinct molecular species. Sedimentation velocity experiments analysis applied to WT Tau (Fig 4) shows that the solution contains 90 % of monomer of Tau with 2.4 S and other forms (10 %) with higher sedimentation coefficient (4S). Because the sedimentation coefficient depends of both the molecular mass and the shape of the studied protein, the 4S sedimentation coefficient could represent a small amount of Tau dimer or a more globular Tau monomer conformation but could not be explained by the presence of Tau large aggregates like pathological fibrils. It is clear that the presence of different peaks in SV experiments explained the polydispersity found in DLS experiments (a major species at more than 80%). If we consider only the major peak in SV we measured a standard sedimentation coefficient (S^0^_20,W_) for WT Tau of 2.4 ± 0.1 S and an estimation of the hydrodynamic radius of 4.5 ± 0.8 nm in agreement with our previous work [28] and with DLS experiments. In this study, we demonstrated that the 16 mutations of serine/ threonine residues into alanine had no impact on the hydrodynamic radius of Tau, irrespective of the experimental methods employed, indicating that the mutant is a suitable model for the study of the two phosphorylation states of Tau. Interestingly, we also found that the two phosphorylation states of AT8 epitope did not modify the hydrodynamic radius of Tau even though its function was described to be modulated by specific phosphorylation patterns [29]. In previous NMR experiments combined with molecular dynamics simulations, Gandhi et al. demonstrated the presence of a kink at residue 207 in pS202/pT205 phosphorylated Tau [8]. The modified Tau protein used in this experiment [8] corresponds to the p-state 1 of the present study. This local conformation was shown to be disrupted by an additional phosphorylation at pS208 corresponding to the p-state 2 phosphorylated Tau [9]. We demonstrated here that these local conformational changes have no effect on the global conformation of Tau. Despres et al. [9] also demonstrated that disruption of the kink in p-state 2 phosphorylated Tau promotes protein aggregation into pathological-like fibers. In the present study, we did not observe fibril formation but we can notice,however, that, in AUC experiments, the area under the very small peak around 4S is larger for p-state 2 (5.5 %) than for p-state1 (2%) (Fig 4 C and D) indicating a propensity of p-state 2 to forms oligomers (dimers) or more globular forms of Tau (Fig 4 C and D). Overall, our experimental observations highlight that the global conformational properties of Tau reflected in its hydrodynamic radius remain unchanged upon phosphorylation on AT8 epitope. To further assess whether computational models reproduce this insensitivity of hydrodynamic radius (Rh) to phosphorylation, we have used the Nygaaard method already applied for unphosphorylated WT Tau protein [17,30]. In their assessment of Rh using the CALVADOS FF, Pesce et al. reported the experimental value of 5.40 +/- 0.20 nm determined by Mukrasch et al. [17,30], However, a re-evaluation of the primary data suggests a value closer to 5.20 +/- 0.20 nm. This adjusted value aligns closely with the linear regression to infinite dilution performed in our study via DLS. Moreover, the experimental flexibility and curvature analysis of Mukrasch et al [17] agree qualitatively with our LCs and LFs for the wild type. We based ourselves on the protocol proposed by Pesce et al. and recovered similar results for the Nygaard Rh despite not using a reweighting technique. This good agreement even with the different p-states tends to confirm the capacity of pCALVADOS to provide relevant Conformational Ensembles for multi-phosphorylated IDPs.

The lack of sensitivity of Rh to mutations, and most importantly to phosphorylations, could be interpreted as a sign that pCALVADOS cannot properly capture the effects of side-chain substitutions due to its coarse-grained nature. Nevertheless, the overall agreement of the calculated Rh with the experimental regression to infinite dilution seems to be telling another story. It is possible that phosphorylation effects are more prominent for intermolecular interactions rather than intramolecular ones, which would explain the dependency on concentration observed on Fig 3 and Table S2. One should however keep in mind the uniqueness problem of CoE, which states that, as bead-like protein models are still complex, there might exist several parameterizations and therefore several CoEs which can fit one target observable [30]. It is therefore possible that pCALVADOS generates ensembles which fit the experimental Rh but are still not biochemically accurate. To evaluate whether this is the case, we must assess whether our CoE fits other experimental properties of multi phosphorylated IDPs.

Our calculations of LCs and LFs revealed that upon multi phosphorylation, the backbone of the Tau mutant locally adopts a less curved and less flexible dynamics (Fig 7). This behavior was previously observed experimentally by Lasorsa et al. on the same system phosphorylated on pSer202 and pThr205 and could be assessed through the analyses of Paramagnetic Relaxation Enhancement effects and the comparison of the relaxation times of the phosphorylated and unphosphorylated forms [19]. This gives weight to the assumption that the CoEs produced by pCALVADOS well describe the experimental observations. LCs and LFs calculated from the simulations suggest that the impact on the average conformation of the backbone does not propagate further than a few residues from the phosphosites (Fig 7). This aligns well with the lack of difference in cw-EPR signals between the different p-states measured on Fig S8. pCALVADOS simulations could thus be a method of choice to guide the selection of positions for nitroxide spin labels, allowing to check multiple positions and whether they stand within the reach of detectable dynamics modification upon phosphorylation.

It is interesting to note that we conducted experiments at low salt concentrations to focus principally on electrostatic interactions. As salt shielding likely impacts the dynamics of multi-phosphorylated Tau in the cell [15], further work is required to mimic the cellular environment. Such investigations could be effectively pursued using Site- Directed Spin Labeling combined with EPR spectroscopy [31] .

Overall, our study constitutes a successful combination of experiments and simulations to study the CoE of a multiphosphorylated IDP at the local and global scales.

## Conclusion

Experimental hydrodynamic radius (Rh) measurements from DLS and AUC for WT Tau and the unphosphorylated as well as progressively phosphorylated (p-state 1 to p-state 2) AT8-phosphorylated mutants were consistent with simulations performed using pCALVADOS 2 forcefield and phospho-residues parameters by Perdikari et al. [14,32]. These findings highlight that specific phosphorylation events at the AT8 epitope do not modulate the hydrodynamic radius and, potentially, the conformational ensemble of Tau.

The generated CoEs display good agreement with experimental Rh using the Nygaard method, as could be expected considering that it is targeted towards IDPs. Our calculations of Local Curvatures and Local Flexibilities confirmed that the mutations on the Tau sequence used to perform selective phosphorylations do not have a significant impact on the local dynamics compared to the wild-type, making the mutant an appropriate surrogate for phosphorylated Tau studies. We also identified a local rigidification and extension of the epitope AT8 proportional to the number of phosphorylation qualitatively supported by several experimental results [17,19]. Long-range effects of phosphorylations were also identified, with an interesting directionality towards the N-terminal end of the Tau protein.

The generation and characterization of the AT8 Tau mutants have laid the groundwork for future studies aimed at elucidating the structure-function relationship of Tau bound on the microtubule. This validated protein will serve as a crucial tool for subsequent studies aimed at establishing a structure-activity relationship for phosphorylations within the AT8 epitope, with a particular focus on residue S208 which was shown to trigger pathological states [9,10]. Although the global size of the mutant remains the same, we show that the phosphorylation of the AT8 epitope still modulates its local conformation. This methodology opens the way to study other pathological phosphorylations in other Tau regions, such as those recognized by the PHF-1 antibody. By systematically investigating the effects of phosphorylation on Tau’s stability, and interactions with microtubules, a perspective of this work is to elucidate the molecular mechanisms underlying the loss of function associated with hyperphosphorylated Tau in neurodegenerative diseases.

## Materials and methods

### WT Tau purification

Tau purification has been already described [33,34] but we will summarize the main step here. The Tau-441 human (hTau40, 2N4R) gene was cloned into a pET-3d vector. This plasmid was transformed into *E. coli* BL21(DE3) cells. The bacteria were grown overnight in a LB medium containing 100 µg/mL ampicillin for antibiotic selection. After dilution, the culture was supplemented with 20 mM glucose to promote growth. The Tau protein production was induced with 0.75 mM isopropyl 1-thio-β-D-galactopyranoside (IPTG) and cells were further incubated for 2 hours 30 min at 37 °C. The cells were harvested by centrifugation at 5000*x g* for 10 mins at 4 °C and then suspended in 2-(N-Morpholino)ethanesulfonic acid 45 mM, Triton X 100 8 mM, DTT 1mM, pH=3.8. The cell suspension was disrupted using a French press, releasing the cellular contents. The lysate was then heated to denature unwanted proteins and clarified by centrifugation at 30000*x g* during 30 min, separating soluble proteins from cell debris. The supernatant containing thermoresistant proteins and Tau was purified using a cation-exchange chromatography column HiTrap SP-HP 5 mL (Cytiva). The column was equilibrated with 45 mM MES, at pH 6.5, and loaded with the supernatant and then a salt gradient was applied. Tau protein bound to the column was eluted with 0.25 M of NaCl. The eluted fractions containing Tau were pooled together. To remove salt excess, the pooled fractions were dialyzed overnight against pure water. The purity of the proteins was verified by SDS-PAGE (Fig S1A). Finally, the Tau protein solution was lyophilized (freeze- dried) for storage.

### Purification and characterization of phosphorylated and unphosphorylated AT8 mutants

The Tau AT8 mutant protein corresponds to a mutated form of Tau 2N4R isoform in which 16 Serine/Threonine were mutated into Alanine to prevent phosphorylation while keeping those of the AT8 phospho-epitope (S202, T205, S208), S198 and T212 [9] (Fig S2). The mutated serine/threonine sites are designed on the basis of the phosphorylation status determined by NMR experiments of hTau40 using recombinant CDK2/cyclin A and GSK3*β* kinases in vitro [9,35]. The mutant was obtained by gene synthesis with optimization of codons for *E. coli* expression and cloned into the pET15b vector (Novagen) into NcoI/XhoI allowing removal of the sequence coding for the polyhistidine tag [35]. ^15^N or ^15^N,^13^C-labeled proteins were produced for functional assays and NMR analyses. Briefly, BL21(DE3) *E. coli* strains transformed with Tau mutants were grown in M9 minimal medium enriched with ^15^N as nitrogen source for uniform isotopic ^15^N-labelling (6g Na_2_HPO_4_, 3g KH_2_PO_4_, 0.5g NaCl, 4g glucose, 1g ^15^NH_4_Cl, 0.5g ^15^N-ISOGRO^®^, 1 mM MgSO_4_, 100 µM CaCl_2_, 100 mg ampicillin per liter) at 37°C. For ^15^N,^13^C-labeling, the medium was prepared the same way with 2g ^13^C_6_- glucose and 0.5g ^15^N,^13^C-ISOGRO^®^. The protein expression was induced by addition of 0.5 mM IPTG for 3 hours. After harvesting the bacteria, the protein was purified by heating the soluble extract at 80°C for 15 minutes followed by cation exchange chromatography (CEx). Then, the CEx fractions were purified by reverse phase HPLC on a Zorbax C8-300SB column (Agilent) using a linear gradient of acetonitrile. The fractions containing the full-length TauAT8 protein were lyophilized, desalted into 20 mM NaPi buffer pH=6.5, dialyzed in water at 4°C overnight, and lyophilized again.

Phosphorylated forms of TauAT8 mutant were produced either by phosphorylation with recombinant CDK2/cyclin A (p-state 1, pS202 + pT205 + pT212) or by a sequential phosphorylation with CDK2/cyclin A and GSK3*β* kinases (p-state 2, pS198 + pS202 + pT205 + pS208 + pT212) as described in [35].

The Tau AT8 mutant and its two phosphorylated forms were analyzed by MALDI-TOF Mass Spectrometry (Axima Assurance, Shimadzu) in a linear positive ion mode with sinapinic acid matrix after ZipTip^®^-C_4_ desalting (Millipore). For the calculation of overall phosphorylation levels, a m/z increment of +80 Da per phosphate group was measured and confirmed by SDS-Page (Figs S2). An NMR analysis was conducted to assess the site-specific phosphorylation sites in each phosphorylation state (Fig S3). Additional mutations were introduced for Site Directed Spin Labeling (SDSL) Electron Paramagnetic Resonance (EPR) experiments as the technique requires the introduction of Cysteine used as target for nitroxide spin labels. Two single cysteine variants of Tau AT8 mutant (variant 1: S185C C291A C322A and variant 2: A227C C291A C322A) were produced as described above and phosphorylated at p-state 1 and 2 prior to spin labeling (Fig 1).

### NMR spectroscopy

#### Sample preparation

For NMR experiments, samples of ^15^N, ^13^C-labeled Tau AT8 (400 μM), and its phosphorylated variants p-state 1 and p-state 2 were prepared in 300 μL of 25 mM phosphate buffer pH 6.4, 50 mM NaCl, 2.5 mM EDTA, 2 mM TCEP in 95% H2O/5% D2O. For chemical shift referencing, 1 mM sodium 3-trimethylsilyl-3,3′,2,2′-d4- propionate was added to the sample and served as the ^1^H reference, with ^15^N and ^13^C chemical shifts indirectly referenced based on ^1^H chemical shifts.

### Data acquisition and processing

5 mm-NMR Shigemi tubes were filled with tau samples for acquisition of 2D ^1^H-^15^N HSQC and 3D experiments at 293 K using a 900 MHz Bruker Avance spectrometer equipped with a triple- resonance cryogenic probe head. Classical backbone assignment was performed on the non-phosphorylated Tau AT8 mutant using various 3D experiments, including HNCACB, HNCO, HNcaCO, HncaNNH, and hNcaNNH, with additional n-CaCON and NCO experiments for proline assignment. All 3D spectra were acquired using non-uniform sampling. Data acquisition and processing were carried out using Bruker Topspin 4.4.0. Following this, NMRFAM-Sparky was employed for NMR data analysis, and the results were cross-validated using the I-PINE web server. The assignment of phosphorylation sites in p-state 1 and p-state 2 Tau AT8 samples has been made through HNCACB and hNcaNNH experiments.

### Determination of WT Tau and Tau AT8 proteins concentration

Lyophilized WT Tau, Tau AT8 mutants and their phosphorylated p-state 1 and p-state 2 forms were resuspended in 20 mM NaPi pH 6.5 buffer containing 1 mM TCEP. Following centrifugation for 10 min at 1500*x g*, the supernatant was diluted to an appropriate concentration for UV-Visible spectroscopy. The absorbance spectra were recorded from 250 to 600 nm using a JASCO-V750 spectrophotometer. To account for light scattering, a mathematical model was employed to correct the absorbance values (y=a+bxe^−c^), where y is the corrected absorbance value, x the measured absorbance value and a, b, and c the fitting constants. Tau concentration was determined using an extinction coefficient of 7700 M^−1^.cm^−1^ at 280 nm with an optical density (OD) between 0.4 and 0.6.

### Protein spin labeling and EPR spectroscopy

Spin labeling variants (positions 185 and 227) of Tau and phosphorylated Tau (p-state 1 and 2) and further continuous wave (cw) EPR analyses were performed as described in [34]. In brief, cysteine reduction was first done by incubating the variants with TCEP (20 mM final, 30 min in ice bath). After removal of TCEP by gel filtration (PD-10 column, *GE Healthcare*, elution buffer 20 mM NaPi pH 6.5), the nitroxide spin label MTSL ([1- oxyl-2,2,5,5-tetramethyl-δ3-pyrroline-3-methyl] methanethiosulfonate; *TRC Inc.*, Toronto, Canada) was added to the sample (10 molar excess, 1 hour in ice bath). Unreacted spin labels were removed by a desalting column PD-10. Cw-EPR spectra were recorded on an Elexsys 500 Bruker spectrometer at X-band (Super High Q sensitivity resonator) using a Bruker N_2_ temperature controller (Bruker ER4131VT). Acquisition parameters were 10mW for microwave power, 0.1 mT and 100 kHz for magnetic field modulation amplitude and frequency respectively.

### Hydrodynamic radius determination by Dynamic Light Scattering (DLS)

Dynamic Light Scattering (DLS) with Malvern Zetasizer Pro equipment was employed to investigate the size distribution and hydrodynamic radius (Rh) of WT Tau, unphosphorylated AT8 mutant and in its phosphorylated p-state 1 and p-state 2 forms. Samples were prepared in a 20 mM NaPi buffer pH 6.5 with 1 mM TCEP and analyzed at 20°C at different protein concentrations (between 0.045 and 1.83 mg/mL). The solvent refractive index and viscosity were set to 1.33 and 1.002 mPa.s, respectively. Data were collected using the Hückel approximation, for triplicates with 150 measurements per sample. The hydrodynamic radius of the particles was determined using the Stokes-Einstein equation: *R*ℎ = *k*_B_*T*/(6*πηD*), where *R*ℎ is the hydrodynamic radius, *k*_B_ is the Boltzmann constant, *T* is the temperature (in K), *η* is the viscosity of the solvent (in Pa.s), and *D* is the diffusion coefficient (in m^2^.s^-1^). The diffusion coefficient was calculated from the measured relaxation time (*τ*) using the following relationship: *τ* = 1/(*q*²*D*), where *q* is the scattering vector.

To determine the distribution of particle sizes, the data were analyzed using two methods: The Hydrodynamic Volume Analysis, providing information about the relative abundance of particles with different sizes; and the Scattered Light Intensity Analysis, providing a more accurate determination of the hydrodynamic radius of the dominant species in the sample.

### Hydrodynamic radius determination by Analytical UltraCentrifugation (AUC)

Sedimentation velocity experiments for WT Tau, Tau AT8 mutants and its phosphorylated p-state 1 and p-state 2 forms were carried out on a Beckman Optima Analytical Ultracentrifuge equipped with absorbance optics. Samples were centrifuged at 60 000 rpm at 20°C into 12 mm double-sector aluminum centerpieces. The absorbance scans were acquired at 275 nm. The raw data were analyzed using the C(S) method implemented in the **SEDFIT** software [36], which fits the sedimentation profiles to a continuous distribution of sedimentation coefficients. This model accounts for diffusion and provides high-resolution separation of species based on their hydrodynamic properties. The C(S) distribution provides the sedimentation coefficient (S_app_) of each species and their relative concentration (in %). To account for the effects of temperature and solvent viscosity, these coefficients were corrected to standard conditions S_app,20,W_ (20°C, in water) using the SEDNTERP software [37] plotted versus protein concentration and extrapolated to infinite dilution to obtain the standard sedimentation coefficient, S^0^_20,W_,. The partial specific volume of WT Tau and Tau mutant was calculated to be 0.736 mL/g and 0.729 mL/g respectively (SEDNTERP software). Finally, to enable a direct comparison across the different Tau phosphorylation states, the distributions were normalized to a uniform concentration.

The solvent density and viscosity were found to be 1.0029 g/mL and 0,01002 poise, respectively. The hydrodynamic radius was obtained using the definition of the sedimentation coefficient and the following Svedberg equation: *R*ℎ = *M*(1 − *<u>υ</u>ρ*) / (6*πηN*_a_*S*). Where *M* is the molecular mass, *<u>υ</u>* is the partial specific volume, ρ is the solvent density, η is the viscosity of the solvent, *N*_a_ the Avogadro’s number and *S* the sedimentation coefficient.

### Statistics

For hydrodynamic radius in DLS experiments and standard sedimentation coefficient S_app,20,W_ determination at infinite dilution, linear regressions were performed. The confidence intervals of Rh and S_app,20,W_ were determined as described below. We first determined the variance of the intercept of the linear regression *b* obtained both in DLS and sedimentation velocity experiments with equation: 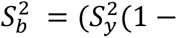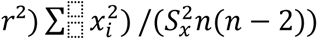.

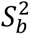 is the variance of the intercept *b*; 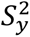 is the variance of the *y* (apparent *R*ℎ or sedimentation coefficient S_app,20,W_); 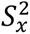 is the variance of the *x* (the protein concentration); *r*² is the coefficient of determination of the linear regression and *n* the number of points. Then, the confidence interval was calculated using equation (3):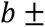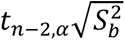. Where *t* is the student parameter obtained for the *n* − 2 degree of freedom and α is the risk with the Bonferroni correction (α = 0.05/number of groups).

### Molecular dynamics simulations: pCALVADOS implementation

The second version of CALVADOS was released in 2022 with improvements regarding dependency on temperature and salt concentration [32] and was recently used on a large-scale study of the Intrinsically Disordered Regions (IDRs) of nearly all the human proteome [38]. Its capacity to generate protein conformational ensembles at low computational cost and its easy implementation in a Google Colab (https://github.com/KULL-Centre/_2023_Tesei_IDRome/blob/main/IDRLab.ipynb) makes CALVADOS a valuable tool to study long IDPs such as Tau. Moreover, since its training set contains the Tau protein and since it has already been used successfully to predict hydrodynamic radii values by Pesce et al. [30], we deemed it a reliable forcefield to use as a base for our study.

We thus performed MD simulations in order to obtain Conformational Ensembles (CoEs) of Tau wild-type and the phosphorylated and unphosphorylated AT8 mutants. At time of writing, CALVADOS did not yet include parameters for phospho-residues, now made available by Rauh et al [13]. We therefore extracted them from the model by Perdikari et al which is also an HPS- based model [14]. This led to the creation of a new parameter set named pCALVADOS, by merging the parameters of CALVADOS 2 [32] with those of doubly charged phosphoserine, phosphothreonine, and phosphotyrosine described by Perdikari *et al* (Table S1 in [14]). Since this kind of “mix-and-match” approach regarding parameters of phospho-residues has proven successful in the past for all-atom force fields, we assumed it might also be the case for coarse-grain approaches, as long as the base model is the same. This assumption was shown to be correct in another work [39], in which a direct comparison with the parameters by Rauh et al showed highly similar dynamic behaviors at global and local scales.

The pCALVADOS parameter set is provided in the following CSV file: https://github.com/Jules-Marien/Articles/blob/main/_2025_Lohberger_Marien_pCALVADOS/pCALVADOS/pCALVADOS_residues_and_phosphoresidues.csv. Phosphoresidues were named based on the Amber nomenclature for the 3-letter denomination and were each given an arbitrary letter as a 1-letter denomination (SEP = B, TPO = J, PTR = O). One simply needs to replace S, T or Y in the sequence by their phospho-residue 1-letter equivalent in order to launch a simulation with phosphorylations. The IDRLab Jupyter Notebook designed by Tesei et al was modified to account for pCALVADOS and to run our simulations of all mutants and p-states. (IDRLab:https://github.com/KULL-Centre/_2023_Tesei_IDRome/blob/main/IDRLab.ipynb). We used OpenMMv7.7 as the simulation engine and simulations were performed in the NVT ensemble with a timestep of 10fs and a Langevin integrator with a 0.01ps^-1^ friction coefficient. The temperature was set at 300K and ionic strength to 0.02 mol.L^⁻1^ to match the DLS experiment. Termini were charged and histidines were kept neutral. Trajectories were run for 100100000 steps, a simulation length sufficient to ensure convergence for an IDP of the length of Tau according to previous studies [32,38]. We also directly tested convergence of global and local observables and showed that it was indeed achieved (see discussion in the Supplementaries, Fig S5 and S9). The 100000 first steps were discarded as equilibration since the protein chain starts as a straight string of beads. Frames were then sampled every 5000 steps, providing a CoE of 20000 evenly-spaced frames.

### From coarse-grain to all-atom

pCALVADOS, as a coarse-grained forcefield, represents each residue as a single bead. It could however be interesting to recover some of the information provided by sidechains, especially as we are interested in phosphorylation. Tesei et al. already proposed a protocol based on the PULCHRA program to convert a CALVADOS trajectory to an all-atom trajectory [40]. PULCHRA is, however, not capable of converting phospho-residues to all-atom representation, and simply provides canonical residues instead. We therefore designed a new protocol to recover all-atom trajectories. Scripts are available at https://github.com/Jules-Marien/Articles/tree/main/_2025_Lohberger_Marien_pCALVADOS/Scripts_CG_2_AA.

At first, PULCHRA was used to generate unphosphorylated all-atom trajectories. A TCL script was then used in combination with VMD to add the proper phosphorylations using CHARMM patches [40] and generate a topology file per conformation. A minimization of 100 steps was performed on each resulting frame using NAMD and the CHARMM36m forcefield, constraining only the backbone heavy atoms to preserve the overall conformation [41,42]. This step allows on one hand to relax any steric and electrostatic tensions induced by adding phosphorylations and also provides a better extrapolation of the sidechains thanks to a state-of-the-art all-atom force field. The relaxed conformations were then re-assembled into an all-atom phosphorylated trajectory. The same protocol was followed for unphosphorylated systems for consistency. Both raw coarse-grain trajectories and relaxed all-atom trajectories are available on Zenodo: https://doi.org/10.5281/zenodo.14900596.

### Rh simulation analysis and new metrics

All analyses were performed using Python scripts based on the MDAnalysis and MDTraj packages [43,44]. Hydrodynamic radii were calculated using the Nygaard method [https://www.sciencedirect.com/science/article/pii/S0006349517306926] as described by Pesce et al. [30], inspiring ourselves from the following scripts: https://github.com/KULL-Centre/papers/tree/main/2022/rh-fwd-model-pesce-et-al/Rh_scripts.

Contact maps were computed by defining a contact as two heavy atoms from two different residues within 5 Å of one another, then calculating the percentage of simulation time during which a pair of residues is in contact. Proteic Menger Curvatures (PMCs), Local Curvatures (LCs) and Local Flexibilities (LFs) were calculated using the previously published Python module [15] :

https://github.com/Jules-Marien/Articles/blob/main/_2024_Marien_nPcollabs/Demo_and_scripts_Proteic_Menger_Curvature/MODULE_Proteic_Menger_Curvature.py. LFs have been shown to correlate significantly with the translational relaxation time T2 for the Tau protein, making the metric a valuable probe of the dynamics of the backbone [39].

Convergence of the simulations is discussed in the Supplementary materials.

## Author contributions

CL, JM and P.B. designed the experiments. CSN and CB designed and prepared the unphosphorylated AT8 mutant and p-state 1 and 2 Tau mutants. CL and PB performed the DLS and UCA experiments and analyzed the data. JM performed all simulations, analyzed the data and modified the CALVADOS 2 IDRLab notebook to incorporate phosphorylation. DA and MT prepared WT Tau. CL performed the determination of the concentration of the mutants. CL and MM performed the EPR experiment and analyzed the data. EB, FXC and CSN performed and analyzed NMR experiments. CL, JM, PB, CSN, CP and SSM participated in the scientific discussions. CL, JM and PB wrote the manuscript. MM and VB supervised the EPR experiments. All authors reviewed and corrected the manuscript.

## Acknowledgement

This study was supported by the French Research Agency ANR-21-CE29-0024 MAGNETAU. This study was supported by the French Research Agency, the French government under the France 2030 investment plan, as part of the Initiative d’Excellence d’Aix-Marseille Université – AMIDEX “AMX-25-HAN-08”, AMX-22-RE-V2-001 and by the LabEx DISTALZ (Development of Innovative Strategies for a Transdisciplinary approach to Alzheimer’s disease, ANR-11-LABX-01). Additional support was provided by the “Initiative d’Excellence” program from the French State (Grant “DYNAMO”, ANR-11-LABX-0011-01 and grant CACSICE, ANR-11-EQPX-0008). The authors are also grateful to the PINT platform facility available at the Institute of NeuroPhysiopathology, to Pavlo Shpak-Kraievskyi and Vecthorus for the accessibility to DLS device, Aix-Marseille Université, CNRS and the NMR and EPR facilities at the French Research Infrastructure INFRANALYTICS (FR2054) and the Aix-Marseille Université EPR center.

## Data availability statement

Raw coarse-grained pCALVADOS trajectories and minimized all-atom trajectories are available in the following Zenodo repository along with scripts allowing the conversion: https://doi.org/10.5281/zenodo.14900596.

pCALVADOS parameters and a jupyter notebook allowing to run a pCALVADOS simulation are available in the following Github repository:https://github.com/Jules-Marien/Articles/tree/main/_2025_Lohberger_Marien_pCALVADOS.

NMR chemical shift assignments of Tau AT8 mutant (from the 2N4R isoform) for 1H, 15N, and 13C backbone including N, HN, Cα, CO, and partial side chain (Cβ) resonances NMR resonances were deposited in the BioMagResBank (BMRB) with entry number 53407.

## Abbreviations

AD: Alzheimer’s disease
AUC: Analytical UltraCentrifugation
CEx: Cation Exchange Chromatography
CoE: Conformational Ensemble
cw: continuous wave
DLS: Dynamic Light Scattering
EPR: Electron Paramagnetic Resonance
HPS: HydroPhobicity Scale
IDP: Intrinsically Disordered Protein
IPTG: isopropyl 1-thio-β-D-galactopyranoside
LC: Local Curvature
LF: Local Flexibility
MD: Molecular Dynamics
MTBD: MicroTubule-Binding Domain
MTSL: [1-oxyl-2,2,5,5-tetramethyl-δ3-pyrroline-3-methyl] methanethiosulfonate
OD: Optical Density
PI: Polydispersity Index
PMC: Proteic Menger Curvature
PRD: Proline Rich Domain
Rh: Hydrodynamic radius
SDSL: Site-Directed Spin Labeling

## Supporting Information

### Convergence of the simulations

**Figure S1: Amino acid sequence of the longest human isoform of Tau (htau40).**

**Figure S2: Phosphorylation level characterization.**

**Table S1 Calculated average and experimental MALDI-TOF masses of WT Tau and the AT8 mutant protein in different phosphorylation states**

**Figure S3: Full views of NMR ^1^H-^15^N HSQC spectra of the unphosphorylated mutant (blue), the mutant in p-state 1 (orange) and in p-state 2 (red).**

**Figure S4: Size distribution by volume (in percentage), correlogram, size distribution by intensity (in percentage) and DLS statistic table of p-state 1 mutant (pSer202 + pThr205 + pThr212) at 9.1 mg/mL.**

**Table S2: Table of DLS data for WT Tau, unphosphorylated AT8 mutant, p-state 1, and p-state 2.**

**Figure S5: Comparison of the LCs (left column) and LFs (right column) calculated on the first half of the trajectory with those calculated on the second half of the trajectory for each residue.**

**Figure S6: Percentages of contacts per residue pair with a maximum of 5% for Tau wild-type(A), the unphosphorylated mutant (B), the mutant in p-state 1 (C) and the mutant in p-state 2 (D).**

**Figure S7: Local Curvatures (A) and Local FLexibilities (B) of the entire Tau monomer.**

**Figure S8: Amplitude normalized cw-EPR spectra recorded at 37°C for the MTSL-labeled Tau at position 185 (A) and position 227 (B).**

**Figure S9: Block-averaging Standard Error (BSE) on the end-to-end distance (Ree, left panels) and the radius of gyration (Rg, right panels) for the pCALVADOS simulations.**

