## Supplementary figures for "Combining experiments and simulations to study the impact of multiple phosphorylations at the AT8 epitope of the Tau protein"

**Running title :** Conformation of phosphorylated AT8-Tau

***Convergence of the simulations***

Convergence of the “global” dynamics of the protein was assessed using the Block-averaging Standard Error (BSE) method described in [1]. We calculated BSE on 2 metrics : Radius of gyration (Rg) and end-to-end distance (Ree).

If the BSE plateaus, we can assume that convergence has been reached, and every observable plateaus for every simulation within a block-size of 10ns. This does not mean that the simulation converges within 10ns, but rather that the decorrelation time scale is of this order. This is a remarkably short time compared to other similar analyses with multi phosphorylated all-atom peptides (see Supplementary Informations of [2][3]) that is probably explained by the very coarse-grained nature of pCALVADOS. The absence of explicit side-chain interactions allows for a faster reorganization of the backbone. We would therefore be wary of the time step scale, as it might actually represent a much longer time than the 1µs that follows from the properties of the model. As can be seen in Figure S4, all BSE tend to plateau except the Rg and Ree of the unphosphorylated mutant which continue to slightly increase up to a block size of 90ns.

Convergence of the “local” dynamics of the protein was assessed by comparing LCs and LFs values computed on the first and second halves of each trajectory (Figure S6). Both observables are observed to have converged since the values by residues display a R2 value of 0.99 in each case, and a RMSE over all residues inferior to 1% of the amplitude of the observables.

These analyses of the convergence of “global” and “local” dynamics show that convergence is reached at all scales for the CALVADOS simulations, a feat which would have cost a drastically larger computational time with an all-atom simulation.

[1] Alan Grossfield and Daniel M. Zuckermann. Quantifying uncertainty and sampling quality in biomolecular simulations.

[2] Marien J, Prévost C, Sacquin-Mora S. n P-Collabs: Investigating Counterion-Mediated Bridges in the Multiply Phosphorylated Tau-R2 Repeat. J Chem Inf Model. 2024;64: 6570–6582. doi:10.1021/acs.jcim.4c00742

[3] Rieloff E, Skepö M. Phosphorylation of a Disordered Peptide-Structural Effects and Force Field Inconsistencies. J Chem Theory Comput. 2020 Mar 10;16(3):1924-1935. doi: 10.1021/acs.jctc.9b01190. Epub 2020 Feb 25. PMID: 32050065.

```
1   MAEPRQEFEV MEDHAGTYGL GDRKDQGGYT MHQDQEGDTD AGLKEAPLQA PTEDGSEEPG   60
61  SETSDAKSAP TAEDVTAPLV DEGAPGKQAA AQPHTEIPEG TTAEEAGIGD TPSLEDEAAG  120
121 HVTQARMVSK SKDGTGSDDK KAKGADGKTK IAAPRGAAPP GQKGQANATR IPAKAPPAPK  180
181 APPSSGEPPK AGDRSGYSAP GSPGTPGSRS RTPSLPAPPT REPKKVAVVR APPKAPSSAK  240
241 SRLQTAPVPM PDLKNVKSKI GATENLKHQP GGGKVQIINK KLDLSNVQSK CGSKDNIKHV  300
301 PGGGSVQIVY KPVDLSKVTS KCGSLGNIHH KPGGGQVEVK SEKLDFKDRV QSKIGALDNI  360
361 THVPGGGNKK IETHKLTFRE NAKAKTDHGA EIVYKAPVVS GDTAPRHLSN VSSTGSIDMV  420
421 DAPQLATLAD EVSASLAKQG L
```

**Figure S1: Amino acid sequence of the longest human isoform of Tau (htau40).** The phosphorylatable Serine/Threonine by CDK2/cyclin A and GSK3$\beta$ kinases mutated in alanine are represented in blue (non-phosphorylatable mutant). In pink the Serine and Threonine phosphorylated by the incubation of Tau with recombinant CDK2/cyclin A (p-state 1: pSer202 + pThr205 + pThr212) and the Serine 198 and 208 phosphorylated in orange with a second incubation of this phosphorylated Tau by GSK3$\beta$ kinases ((p-state 2: pSer198 + pSer202 + pThr205 +pSer208 + pThr212). In purple the single Cysteine variant at positions 185 (non-phosphorylatable, p-state 1 or p-state 2 S185C). In light green the single Cysteine variant at position 227 (non-phosphorylatable, p-state 1 or p-state 2 A227C). In these two variants the 2 natural cysteines in position 291 and 322 were mutated in Alanines.

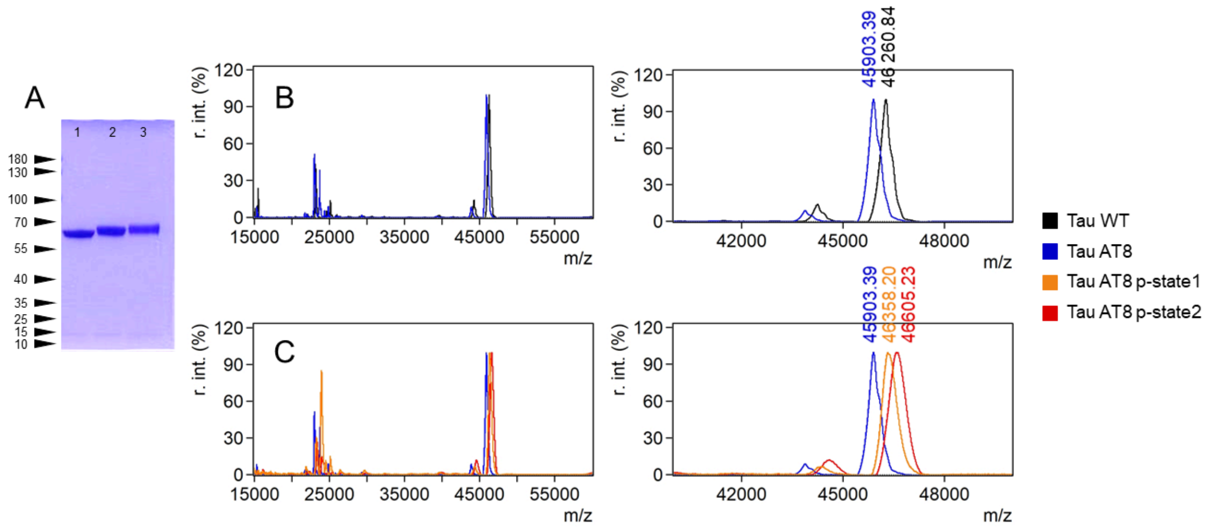


**Figure S2: Phosphorylation level characterization A.** SDS-PAGE of non-phosphorylated mutant (lane1), p-state 1 mutant (lane 2) and p-state 2 mutant (lane 3) deposit of 4 µg per lane. **B, C**. MALDI-TOF mass spectrometry analysis: full *m/z* spectra (left panels) and expanded views at *z=1* (right panels) of Tau proteins are depicted. Annotated signals correspond to the full-length Tau proteins (wild-type or mutant). A small proportion of a Tau fragment corresponding to protein truncation during the purification step is shown at a *m/z* ratio below 45,000. Panels B show the comparison of unphosphorylated wild-type (in black) and mutant Tau (in blue). Panels C show the unphosphorylated mutant (in blue), the mutant in p-state 1 (in orange) and the mutant in p-state 2 (in red), highlighting the differences of *m/z* ratios for the different proteoforms of Tau mutant.

| Construct | Calc. average mass (15N-enriched, Da) | Exp. mass (Da) | Δ (Da) calc. – exp | FWHM/ Resolution | Δ (vs. previous state, Da) | Equivalent phosphorylation increase* |
|---|---|---|---|---|---|---|
| WT tau | 46,426.51 | 46,260.84 | -165.67 | 356.756/ 130 | – | – |
| AT8 mutant | 46,072.33 | 45,903.39 | -168.94 | 322.201/ 142 | -357.45 (vs. WT) | – |
| P-state 1 (CDK2) | – | 46,358.20 | | 439.326/ 106 | +454.81 (vs. AT8) | +5.68 $PO_3$ |
| P-state 2 (CDK2 + GSK3β) | – | 46,605.23 | | 507.992/ 92 | +247.03 (vs. P-state 1) | +3.09 $PO_3$ |

**<u>Table S1</u>** : Calculated average and experimental MALDI-TOF masses of WT tau and the AT8 mutant protein in different phosphorylation states. Average theoretical masses were calculated for uniformly ^15N-enriched proteins. Experimental masses were determined by MALDI-TOF mass spectrometry. The difference between calculated and experimental masses (Δ) is reported for the non-phosphorylated proteins. For phosphorylated species, mass increases relative to the preceding phosphorylation state are shown. Peak widths are reported as full width at half maximum (FWHM), together with the corresponding mass resolution (m/z/FWHM). The equivalent phosphorylation increase was estimated using a mass increment of 79.97 Da per phosphate group and is provided for qualitative interpretation only, as the limited resolution of the intact protein MALDI-TOF measurements does not allow accurate determination of the exact number or occupancy of phosphorylation sites.

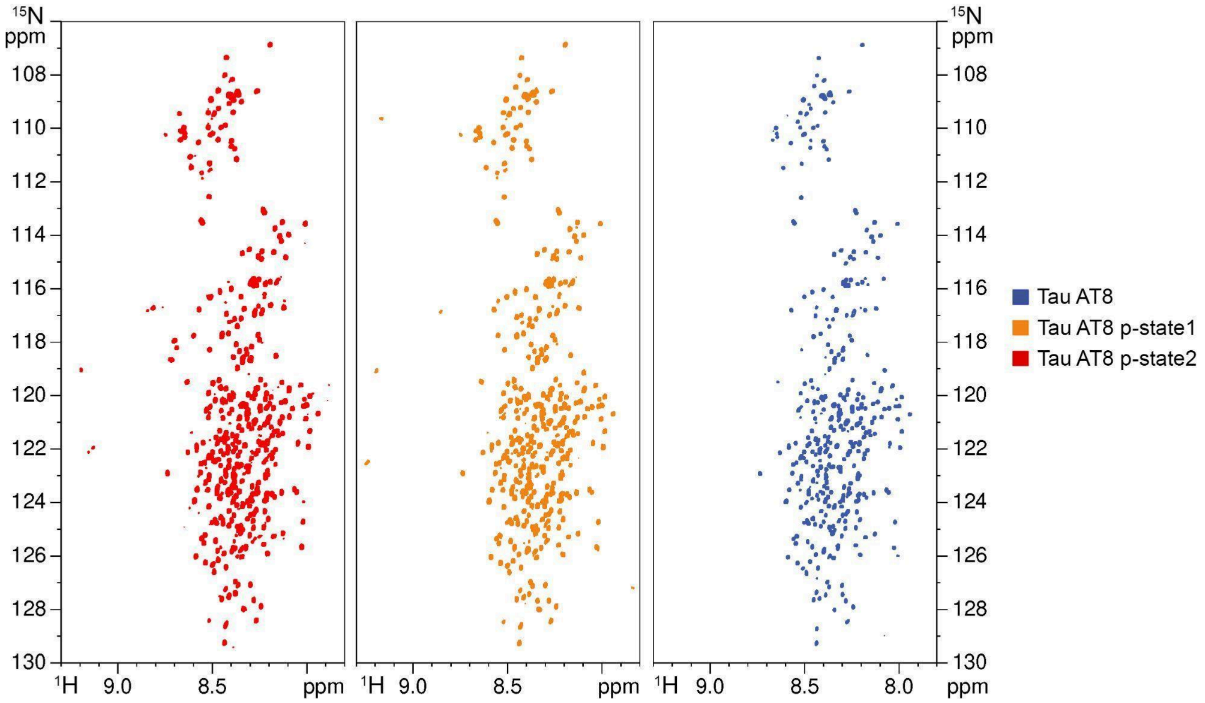


**Figure S3: Full views of NMR $^{1}H$-$^{15}N$ HSQC spectra of the unphosphorylated mutant (blue), the mutant in p-state 1 (orange) and in p-state 2 (red).**

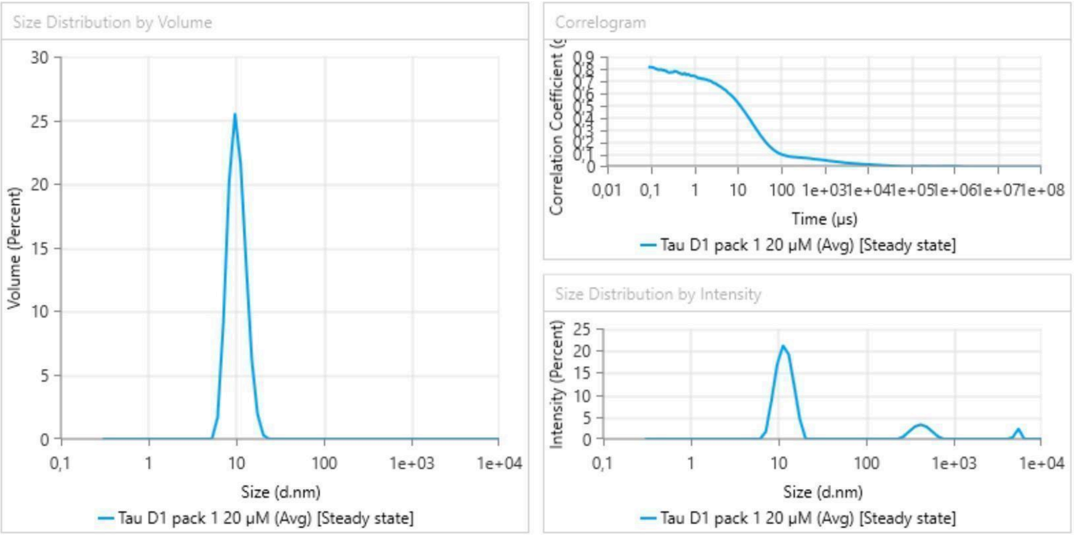


| Name | Mean | Standard Deviation | RSD | Minimum | Maximum |
|---|---|---|---|---|---|
| Z-Average (nm) | 57,95 | - | - | 57,95 | 57,95 |
| Polydispersity Index (PI) | 0,2405 | - | - | 0,2405 | 0,2405 |
| Peak 1 Mean by Intensity ordered by area (nm) | 11,85 | - | - | 11,85 | 11,85 |
| Peak 1 Area by Intensity ordered by area (%) | 83,81 | - | - | 83,81 | 83,81 |
| Peak 2 Mean by Intensity ordered by area (nm) | 430,3 | - | - | 430,3 | 430,3 |
| Peak 2 Area by Intensity ordered by area (%) | 13,49 | - | - | 13,49 | 13,49 |

**Figure S4: Size distribution by volume (in percentage), correlogram, size distribution by intensity (in percentage) and DLS statistic table of p-state 1 mutant (pSer202 + pThr205 + pThr212) at 0.91 mg/mL.** Data are an average of triplicates. Volume (in percentage) as a function of the hydrodynamic diameter (nm), correlation coefficient (g2-1) as a function of time (µs) and Intensity (percentage) as a function of the hydrodynamic diameter (nm).

| WT Tau | | | | | | | | |
|---|---|---|---|---|---|---|---|---|
| **[mg/mL]** | 0.092 | 0.23 | 0.37 | 0.46 | 0.55 | 0.92 | 1.46 | 1.83 |
| **Rh (nm)** | 4.7 | 5.14 | 5.35 | 5.42 | 5.03 | 5.66 | 4.98 | 5.93 |
| **SD** | 1.8 | 0.8 | 0.3 | 0.5 | 0.1 | 0.07 | 0.3 | 0.7 |

| unphosphorylated AT8 mutant | | | | | | | | | |
|---|---|---|---|---|---|---|---|---|---|
| **[mg/mL]** | 0.045 | 0.091 | 0.23 | 0.36 | 0.45 | 0.68 | 0.91 | 1.14 | 1.45 |
| **Rh (nm)** | 5.51 | 4.83 | 4.73 | 5.3 | 5.74 | 6.03 | 5.66 | 6.13 | 6.78 |
| **SD** | 0.7 | 0.4 | 1.4 | 0.1 | 0.6 | 0.5 | 0.6 | 0.2 | 0.3 |

| p-state 1 | | | | | | | | | |
|---|---|---|---|---|---|---|---|---|---|
| **[mg/mL]** | 0.091 | 0.23 | 0.36 | 0.54 | 0.68 | 0.82 | 0.91 | 1.14 | 1.45 |
| **Rh (nm)** | 5.69 | 5.67 | 6.35 | 5.95 | 5.92 | 6.12 | 5.92 | 5.8 | 6.24 |
| **SD** | 0.7 | 0.7 | 0.5 | 0.3 | 0.8 | 0.3 | 0.5 | 1.8 | 0.01 |

| p-state 2 | | | | | | | | | | |
|---|---|---|---|---|---|---|---|---|---|---|
| **[mg/mL]** | 0.045 | 0.091 | 0.23 | 0.36 | 0.45 | 0.54 | 0.68 | 0.91 | 1.14 | 1.45 |
| **Rh (nm)** | 5.46 | 4.59 | 5.16 | 5.62 | 6.01 | 5.73 | 6.47 | 5.95 | 6.19 | 7.19 |
| **SD** | 1.3 | 1.1 | 0.17 | 0.2 | 0.4 | 0.2 | 0.6 | 0.4 | 0.7 | 0.2 |

**Table S2: Table of DLS data for WT Tau, unphosphorylated AT8 mutant, p-state 1, and p-state 2.** Concentrations in mg/mL, hydrodynamic radius in nm, and Standard Deviation (average of triplicates).

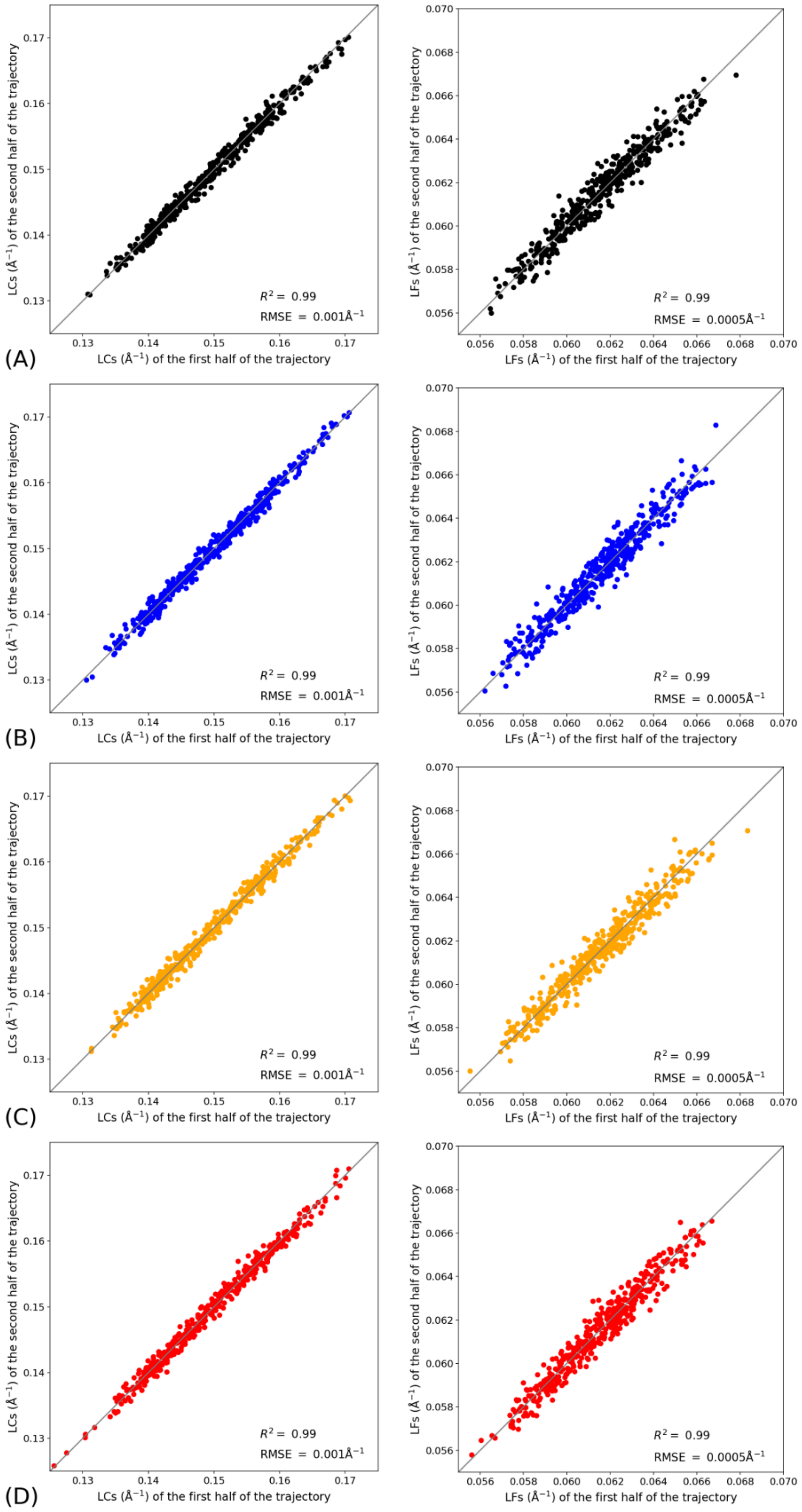


**Figure S5: Comparison of the LCs (left column) and LFs (right column) calculated on the first half of the trajectory with those calculated on the second half of the trajectory for each residue.** A) wild-type sequence. B) Unphosphorylated mutant. C) Mutant in p-state 1. D) Mutant in p-state 2.

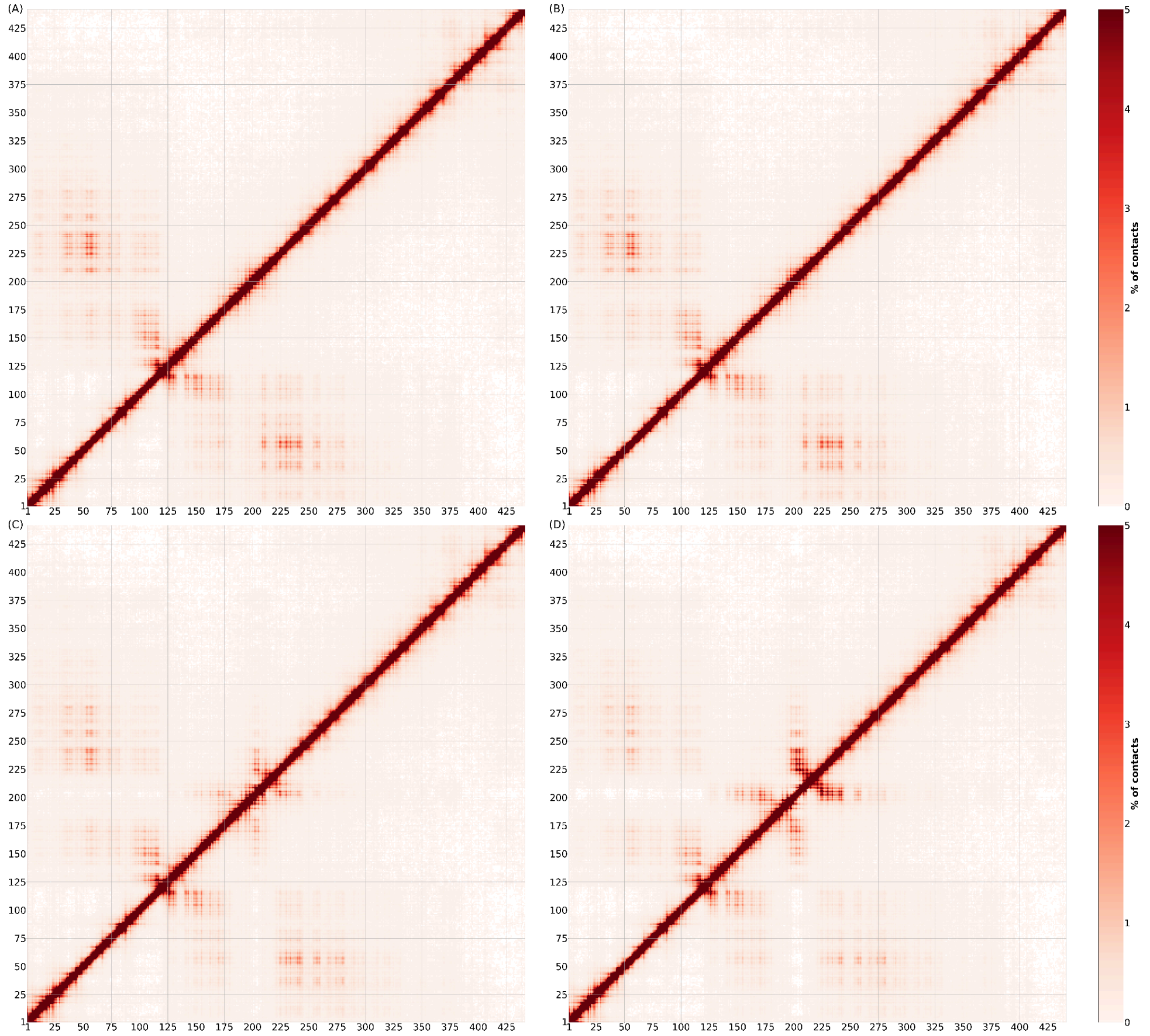


**Figure S6: Percentages of contacts per residue pair with a maximum of 5% for Tau wild-type(A), the unphosphorylated mutant (B), the mutant in p-state 1 (C ) and the mutant in p-state 2 (D).** Contacts run through light red to deep red, absence of contacts is in white.

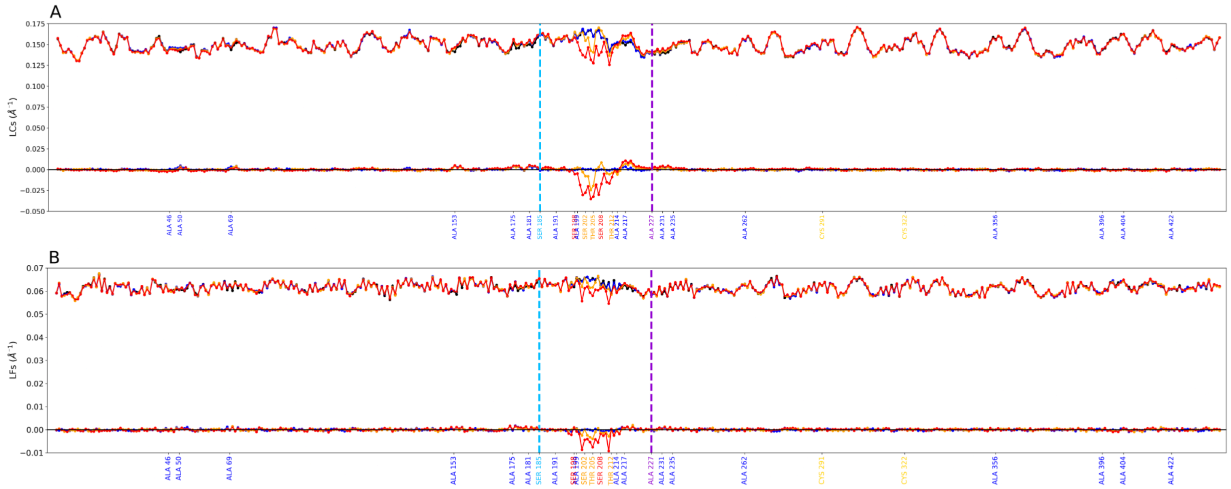


**Figure S7: Local Curvatures (A) and Local FLexibilities (B) of the entire Tau monomer.** Upper curves on each graph represent Local Curvatures (A) and Local Flexibilities (B) along the sequence for wild-type Tau (black), the unphosphorylated mutant (blue), the mutant in p-state 1 (orange) and the mutant in p-state 2 (red) The lower curves on each graph represent the difference to the wild-type. Positive values represent a gain in curvature/flexibility respectively and negative values a loss. Residues in blue have been mutated to alanines compared to Tau wild-type. Residues in orange are phosphorylated in p-state 1, residues in red in p-states 1 and 2. The superposed residues are SER198 and ALA199. The mutated sites for cw-EPR experiments S185C and A227C are indicated in blue and purple dashed lines respectively.

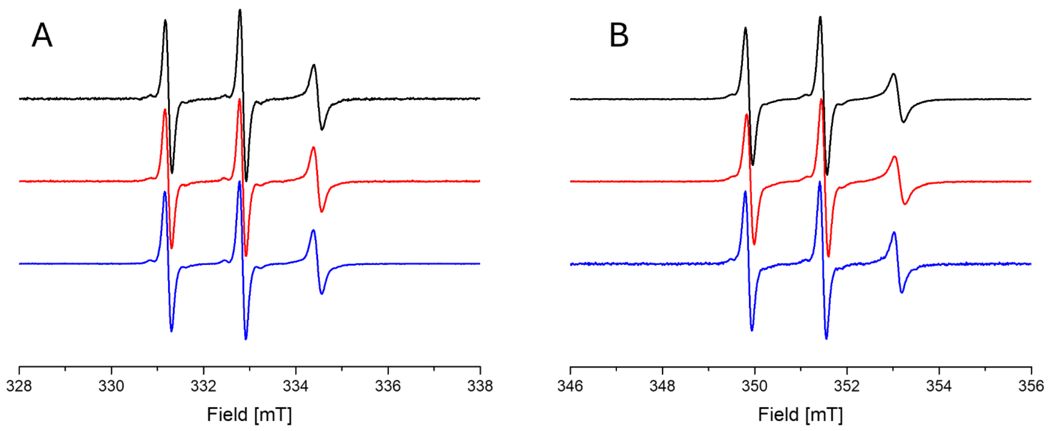


**Figure S8: Amplitude normalized cw-EPR spectra recorded at 37°C for the MTSL-labeled Tau at position 185 (A) and position 227 (B).** Spectra obtained for the unphosphorylated Tau mutants (black), for the p-state 1 (red) and for the p-state 2 (blue) are displayed. The three narrow lines indicate a very high mobility of the nitroxide spin label, a shape typical for IDPs. No significant difference is observed between the two phosphorylated states and the unphosphorylated one, revealing that the local flexibility around aa 185 (downstream the AT8 region) and 227 (upstream the AT8 region) is not modified from each side of the targeted phosphorylated region.

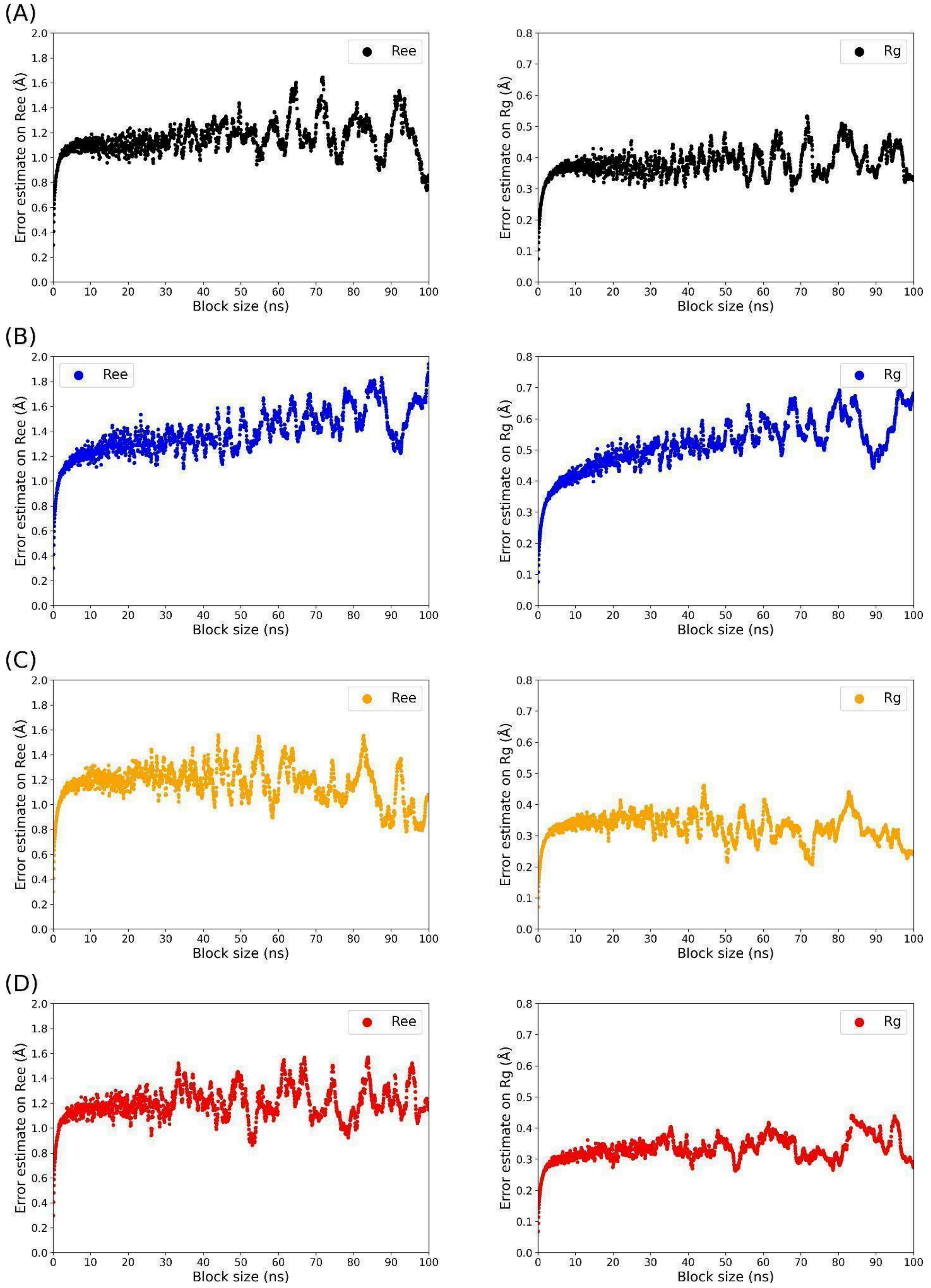


**Figure S9: Block-averaging Standard Error (BSE) on the end-to-end distance (Ree, left panels) and the radius of gyration (Rg, right panels) for the pCALVADOS simulations.** BSEs were calculated for the wild-type sequence (A), the unphosphorylated mutant (B), the mutant in p-state 1 (C) and the mutant in p-state 2 (D).
